# Single-cell transcriptome of developing inhibitory neurons reveals expanding and contracting modes of diversification

**DOI:** 10.1101/2025.02.19.636192

**Authors:** Minhui Liu, Facundo Ferrero Restelli, Elia Micoli, Giulia Barbiera, Rani Moors, Evelien Nouboers, Malou Reverendo, Jessica Xinyun Du, Hannah Bertels, Dimitris Konstantopoulos, Keimpe Wierd, Aya Takeoka, Giordano Lippi, Lynette Lim

## Abstract

The cerebral cortex relies on vastly different types of inhibitory neurons to compute. How this diversity emerges during development remains an open question. The rarity of individual inhibitory neuron types often leads to their underrepresentation in single-cell RNA sequencing (scRNAseq) datasets, limiting insights into their developmental trajectories. To address this problem, we developed a computational pipeline to enrich and integrate rare cell types across multiple datasets. Applying this approach to somatostatin-expressing (SST+) inhibitory neurons—the most diverse inhibitory cell class in the cortex—we constructed the transcriptomic maps, Dev-SST-v1 and Dev-SST-v2, a comprehensive resource containing mouse scRNAseq data of over 51,000 SST+ neurons. We identify three principal groups—Martinotti cells (MCs), non-Martinotti cells (nMCs), and long-range projecting neurons (LRPs)—each following distinct diversification trajectories. MCs commit early, with distinct embryonic and neonatal clusters that map directly to adult counterparts. In contrast, nMCs diversify gradually, with each developmental cluster giving rise to multiple adult cell types. LRPs follow a unique ‘contracting’ mode. Initially, two clusters are present until postnatal day 5 (P5), but by P7, one type is eliminated through programmed cell death, leaving a single surviving population. This transient LRP type is also found in the fetal human cortex, revealing an evolutionarily conserved feature of cortical development. Together, these findings highlight three distinct modes of SST+ neuronal diversification—invariant, expanding, and contracting—offering a new framework to understand how the large repertoire of inhibitory neurons emerges during development.

## Introduction

Cortical inhibitory neurons orchestrate information processing, coordinate network synchrony and are essential to circuit homeostasis ^1–7^. Dysregulation in their development has been implicated in a range of neurological disorders ^7–13^, including epilepsy, schizophrenia, and autism. This underscores the critical need to understand how these neurons develop and diversify.

Advances in single-cell and single-nucleus RNA sequencing (sc/snRNA-seq) have transformed the study of brain tissue, which contains heterogeneous cell types. However, adequate representation of cortical inhibitory neurons in scRNAseq datasets remains particularly challenging due to their rarity and their diversity. Resolving their developmental trajectories requires either extensive enrichment or sequencing of large populations—approaches that are not standard in most experiments.

In the adult mouse cortex and hippocampus, inhibitory neurons make up only ∼15% of cortical neurons, with the remaining ∼85% being excitatory glutamatergic neurons ^4,7,14^. Within this minority population, SST+ neurons stand out due to their exceptionally high transcriptional and functional diversity ^15–19^, further contributing to poor representation of specific cell types. Recent efforts to construct large-scale single-cell atlases, such as the ABC Atlas ^15,20^, have introduced hierarchical classification systems—class, subclass, supertype, and type or clusters—that improve the granularity of cell type annotation. However, rare cell types remain severely underrepresented; for instance, 16% of SST+ types (or clusters) are represented by fewer than 150 cells, compared to only 2.7% of excitatory pyramidal neuron types ^15,20^. Similar underrepresentation persists in human brain scRNA-seq datasets, where rare inhibitory neuron types, such as Sst-Chodl neurons, are particularly sparse ^21^.

The challenge of poor representation is further compounded in developmental datasets ^22–26^. During early development, transcriptional profiles are dominated by immature signatures, which obscure cellular diversity ^22–26^. For SST+ neurons, encompassing Martinotti cells (MCs), non-Martinotti cells (nMCs), and long-range projecting neurons (LRPs) ^4,6,7,23,26–30^, understanding developmental trajectories is particularly difficult due to the lack of fine-scale resolution in existing data.

To overcome these limitations, we developed a computational pipeline to enhance the detection and integration of rare neuronal cell types in developmental scRNA-seq datasets. Our computational pipeline consists of three modules: (i) rare cell extraction, (ii) dataset integrability assessment, and (iii) hierarchical clustering for atlas construction. Using SST+ neurons as a model system, we constructed two RNAmaps we named as Dev-SST-v1 and Dev-SST-v2, high-resolution developmental resources that span embryonic day (E)16.5 to postnatal day 7 (P10). Both atlases include over 50,000 SST+ neurons, representing a seven-fold increase in sampling compared to our previous efforts ^23^.

Our analysis reveals three principal groups of SST+ neurons—Martinotti cells (MCs), non-Martinotti cells (nMCs), and long-range projecting neurons (LRPs)—each with distinct diversification and developmental trajectories. MCs follow an invariant trajectory, where early developmental clusters align closely with adult cell types, similar to excitatory pyramidal neurons ^31^, Chandelier, and Meis2+ interneurons ^22,32^. nMCs exhibit an expanding trajectory, where a single neonatal cluster gives rise to multiple adult types, resembling retinal ganglion cell development ^33^. LRPs undergo a “contracting” trajectory, with two early postnatal clusters persisting until P5, after which one is selectively eliminated through programmed cell death, leaving a single surviving type in adulthood.

In summary, our computational pipeline is a powerful resource for studying rare neuronal cell types. Dev-SST-v1 and v2, the most comprehensive single cell inhibitory neuron atlas to date, revealed an unprecedented palette of cell diversification strategies in the developing mammalian cortex.

## Results

### Developing somatostatin-expressing inhibitory neurons are underrepresented in most scRNAseq datasets

Somatostatin-expressing (SST+) inhibitory neurons represent about 3-4% of all cortical neurons, and are one of the most diverse classes among inhibitory neurons ^7,15–17,19,34^. In adult mouse neocortex, scRNAseq datasets have shown that these inhibitory cells can be grouped into 19 supertypes and 45 to 73 types or clusters ^15,17,20^. To investigate the development of these rare neurons, we constructed a high-resolution, integrable single-cell atlas by extracting SST+ cells from multiple datasets. Our computational pipeline consists of three modules designed for this purpose (**Fig. 1a**).

**Figure 1.**
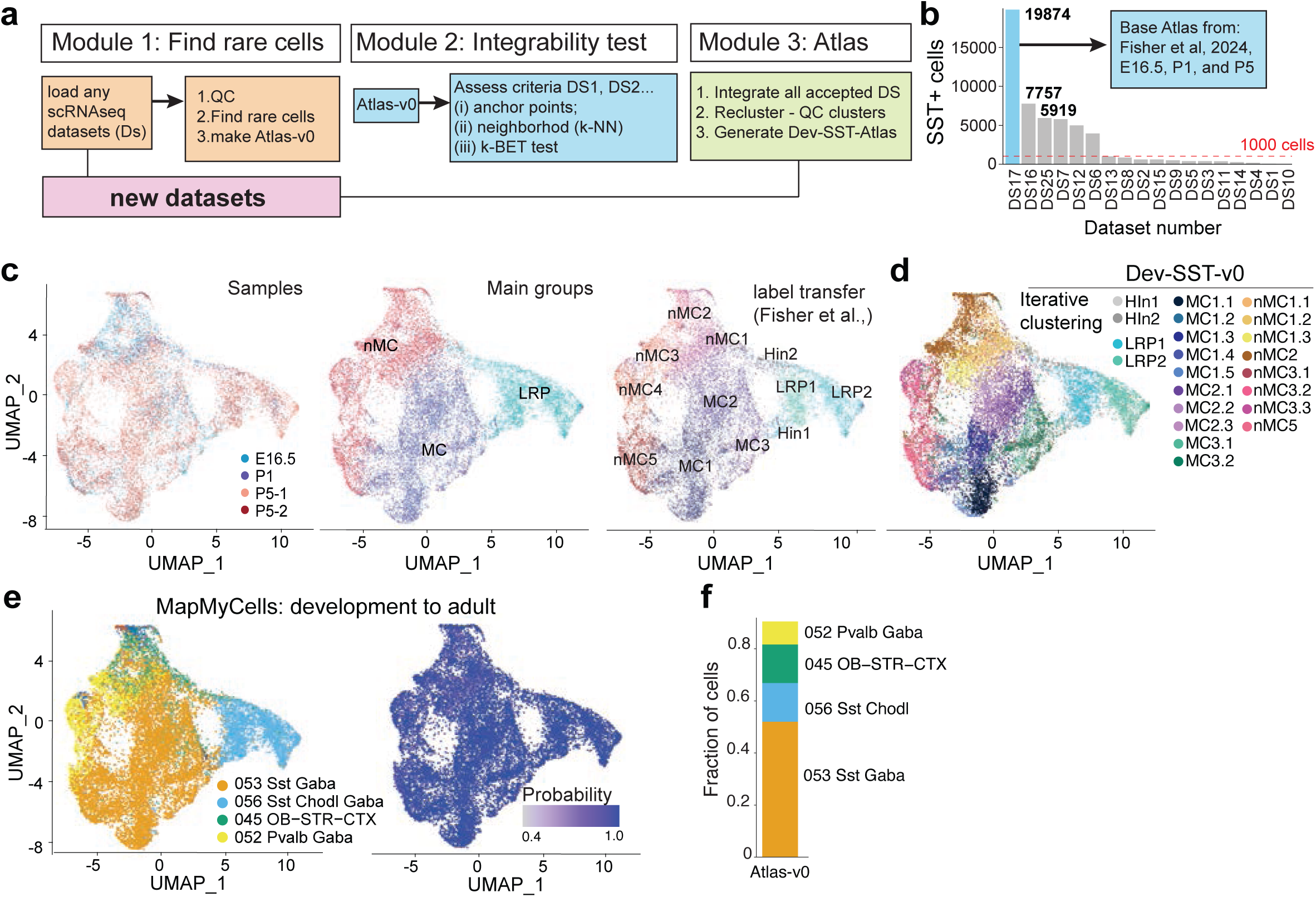
Integrative scRNAseq developmental atlas for SST+ inhibitory neurons. (**a**) Schematic of the three-step integrative pipeline for generating an atlas. Step 1: Selection of rare cell classes (SST+) from publicly available scRNAseq datasets. Step 2: Assessment of dataset integrability using anchor points, k-nearest-neighbor (k-NN) graph evaluation, and k-BET testing. Step 3: Construction of the new developmental SST+ atlas (Dev-SST-Atlas) by integrating accepted datasets, reclustering, and quality control. (**b**) Bar plot showing the distribution of SST+ cells identified in each dataset in Module 1, highlighting variability in SST+ cell representation across datasets. (**c)** UMAP visualization of Dev-SST-v0, comprising 19,874 SST+ cells, annotated by sample types, main groups, and subtype labels transferred from Fisher et al. (2024). (**d**) UMAP visualization of Dev-SST-v0 annotated by iterative clustering parameters set in this paper. (**e**) UMAP embedding of Dev-SST-v0 annotated using MapMyCells with ABC Atlas (Yao et al. 2023)^15^, overlaying mapping probabilities for each cell. (**f**) Stacked bar plot depicting the relative abundance of subclasses in Atlas-v0, restricted to cells with mapping high probabilities (>0.9).

In the first module (Module 1), we systematically survey publicly available scRNAseq datasets from the developing mouse cortex spanning embryonic day (E)12.5 to postnatal day (P)10 (**Extended Data Table 1**), collectively comprising more than 1.5 million cells ^23,26,35–49^. Initial quality control measures included filtering out cells with low counts and high mitochondrial DNA content (Methods; **Extended Data Fig. 1a**). SST+ cells were identified by clustering each dataset and selecting populations expressing both *Sst* and *Lhx6*, two key markers of developing inhibitory neurons ^7,24,27,50,51^.To exclude contaminating interneurons, we removed Meis2+ (SST-negative) cells known to appear in *Sst^cre^*driver lines ^23^.

Of the 18 datasets analyzed (DS1–DS17, DS25 ^23,26,35–49)^, only seven contained more than 1,000 SST+ neurons (**Extended Data Table 1a**, **Fig. 1b**). The largest dataset, from Fisher et al. (2024) ^23^, contained ∼19,000 SST+ cells and was used to generate a preliminary reference, Atlas-v0 or Dev-SST-v0, incorporating data from four experiments spanning three developmental stages (E16.5, P1, P5).

To annotate Dev-SST-v0, we performed label transfer using annotations from Fisher et al. ^23^ (**Fig 1c**). Iterative clustering of Dev-SST-v0 (see methods, **Extended Data Fig 1a-c**) reveal more clusters than label transfer (**Fig 1d**). As a control, we use the same clustering parameter (see methods) and applied to the original datasets from Fisher et al., (2024), and found that our current parameter achieved a similar number of clusters (**Extended Data Fig 2a, 2b)**. For each dataset, we subset 1000 cells and compute the mean anchor point against the 3 other datasets (**Extended Data Fig 2c, 2d)**, and computed the anchor distribution among the four reference samples, which range from 1021 to 2775 (**Extended Data Fig 2d)**.

Then, we mapped developmental cells onto adult inhibitory neuron reference data from Yao et al. (2023) ^15^, using the Allen Institute’s *MapMyCells* algorithm (RRID:SCR_024672). At the subclass level, more than 75% of Dev-SST-v0 cells aligned with adult SST+ inhibitory neurons (Sst 053, Sst-Chodl 056; **Fig. 1e**, **1f**), with high confidence (>0.9 probability). Approximately 15% of cells mapped to Pvalb (052) or olfactory-striatal inhibitory neurons (045 OB-STR; **Fig. 1e**). Given that SST+ cells in Dev-SST-v0 were enriched using fluorescence-activated cell sorting (FACS) from *Sst^cre^;RCE* animals, which contain minimal (<1%) Pvalb+ cells ^23^, this suggests that a subset of developing SST+ neurons may transcriptomically resemble adult Pvalb+ and olfactory-striatal interneurons. Dev-SST-v0 has 4 distinct datasets. Overall, Module 1 of our computational pipeline selectively and accurately extracts rare cell types.

### A generalized computational pipeline to assess integrability of different datasets

The second module (Module 2) of our pipeline was designed to examine structural similarities between two distinct datasets (**Fig. 1a, Extended Data Fig. 1b**). Since only 7 datasets contain more than 1000 SST+ (**Extended Data Table 1**), we reasoned these small numbers of cells would not give us sufficient representation to build a high resolution atlas for SST+ neurons. Therefore, we generated 8 more datasets (DS18 to DS24, DS26, **Extended Data Table 1**), all containing more than 1000 SST+ cells from the developing mouse cortex from E16.5 to P5 (**Fig. 2a**). Most of these are enriched by FACS as previously described ^23^ (see methods, and **Extended Data Table 1**). For these unpublished datasets, we applied the computational pipeline of Module 1 to extract all SST+ cells **(Fig. 2a)**.

**Figure 2.**
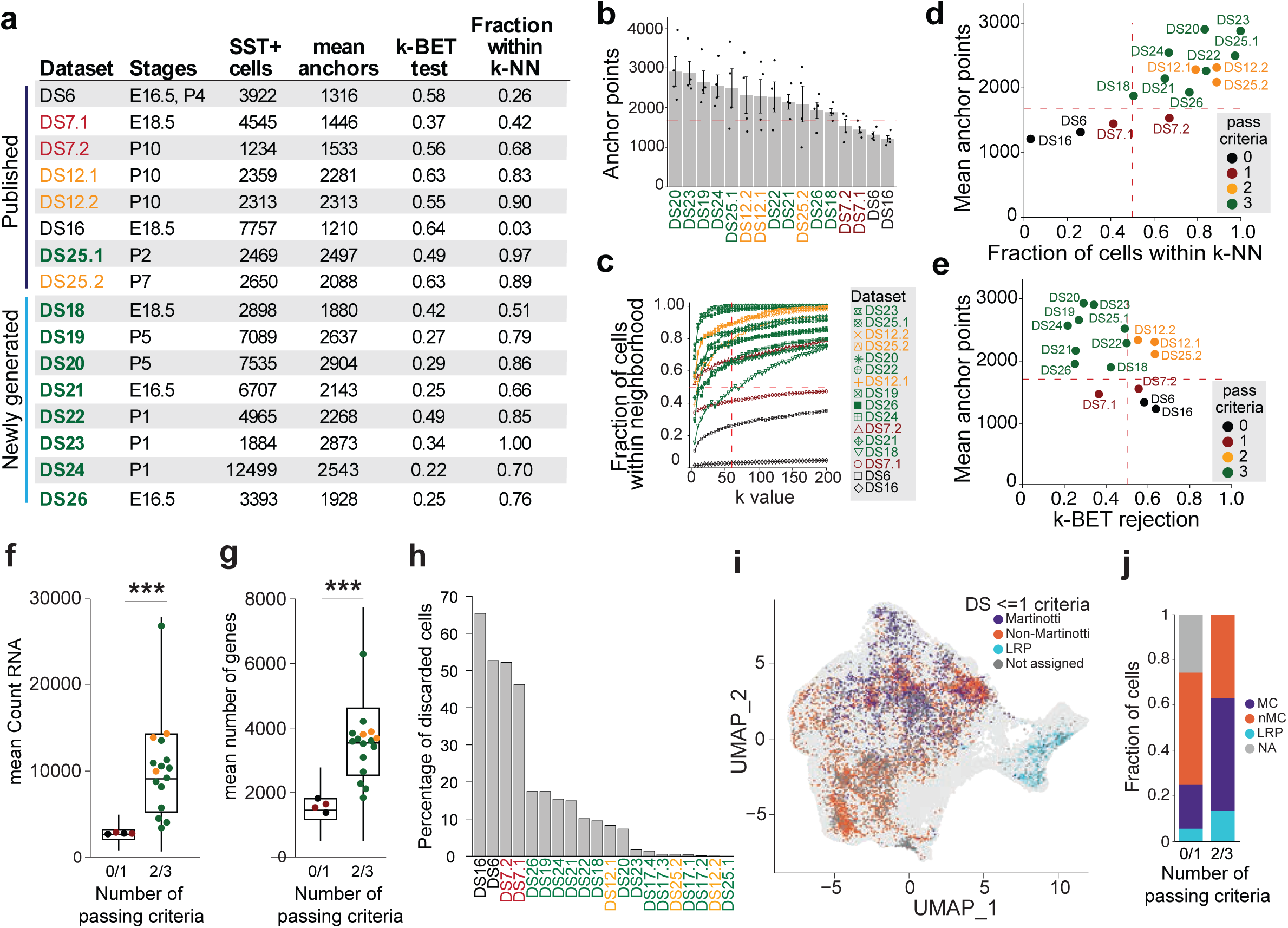
Computational pipeline to assess the integrability of different datasets. (**a**) Tabular overview of datasets used to test integrability with Dev-SST-v0, including both newly generated and published datasets. The table lists SST+ cell counts, normalized mean anchor points, k-BET (k-nearest-neighbor batch effect test) rejection rate, and fractions of cells within Dev-SST-v0 (k-NN, k = 60) for each dataset. (**b**) Bar plot of normalized mean anchor points calculated using Seurat’s CCA integration function between each dataset (subsetting randomly 1000 cells) and Dev-SST-v0. Datasets with >1686 mean anchor points (horizontal dashed line) were considered integrable. (**c**) Fraction of cells in each dataset within Dev-SST-v0 neighborhoods with increasing k value. The dashed line (k = 60) indicates the acceptance threshold of >50%. (**d**, **e**) Dot plot of (d) fraction cells within k-NN (k=60) and (e) k-BET rejection plotted against anchor points rates for each dataset. Datasets passing the three integrability criteria (in green) have mean anchor points >1686, fraction within k-NN >50%, and k-BET rejection rate <0.5. **(f)** Box plot of mean count of RNA from datasets assessed passing either 0/1 or 2/3 criteria, ***p<0.001. (**h**) Percentage of discarded cells post-QC from each datasets. (**i**) UMAP visualization of all DS and highlight of DS passing 0/1 criteria. (**j**) Stacked bar plot of cells assigned as MC, nMC, LRP, or not assigned in DS passing 0/1 pr 2/3 criteria.

In stages earlier than E16.5, such as E13.5, we noted that only few SST+ cells have migrated to the cortex (**Extended Data Fig 2e, 2f).** Isolation of MGE at E12.5 captures newly born SST+ interneurons (**Extended Data Fig 2g, h, i).** While these post-mitotic neurons could be mapped to large groups such as MC/nMC/LRP with similar distribution as later timepoints (**Extended Data Fig 2i)**, the mapping probability of each cluster is much poorer than samples from E16.5 onwards (**Extended Data Fig 2j)**. Furthermore, post E16.5, we do not detect differences in mapping probability between clusters of MC or nMC (**Extended Data Fig 2k)**. Therefore, we focused our subsequent analyses with samples that are E16.5 and older.

Next, we proceeded to assess if the SST+ inhibitory neurons from different sources, both published and newly generated datasets, can be integrated using three distinct criteria. These criteria were designed to capture complementary aspects of dataset compatibility — transcriptomic similarity, local structural consistency, and batch mixing quality. For the first criterion, we quantified the number of anchor points by randomly subsampling 1,000 cells per dataset (see Methods) and applying the Seurat CCA integration between the SST⁺ cells from each dataset and those in Dev-SST-v0. Because Dev-SST-v0 includes four distinct reference samples from Fisher et al. ^23^, anchor points were computed for each new dataset relative to each reference sample (see Methods). We set the acceptance threshold at datasets with >1,686 normalised mean anchor points (**Fig. 2a, 2b**), corresponding to the 25th percentile of the anchor distribution among the four reference samples in Dev-SST-v0 (**Extended Data Fig. 2d**).

For the second criteria, we determined the k-nearest neighbor (k-NN) in each dataset using principal components computed on the integrated counts matrix. Then, we evaluated the neighborhood composition for all the cells in each dataset, and determined the percentage of cells that have similar neighborhoods (k-NN), which is 50% in Atlas-v0. By adjusting the neighborhood size (k from 0 to 200), we estimated the structure similarly of each dataset to Atlas-v0 (**Fig. 2c**). We set the acceptance criteria for each dataset as having >50% of cells within the same k-NN (k=60), as observed in Atlas-v0 (**Fig. 2d**).

For the last criterion, we calculated the k-BET rejection rate ^52^, a test commonly used for assessing samples similarly from different batches. We set the acceptance criteria as k-BET < 0.5 (**Fig. 2a, 2e**). Among the 13 datasets subjected to these tests (**Fig. 2a, 2d, 2e**), 7 datasets passed all three criteria and they were used to generate a developmental atlas for SST+ neurons in Module 3.

To evaluate whether our integration criteria reflected true dataset quality, we compared several transcriptomic and clustering metrics between datasets that passed ≤1 criterion and those passing ≥2 criteria. Datasets meeting ≥2 criteria exhibited statistically higher mean RNA counts and mean gene numbers per cell (**Fig. 2f, 2g, \*\*\***p<0.001) metrics that reflect sequencing depth and transcript detection efficiency. They also showed a lower proportion of discarded cells during quality control (<20%), whereas datasets passing ≤1 criterion had a higher rate of cell exclusion (>40%) (**Fig. 2h**), indicating that integrability is closely linked to overall cell quality. Among the retained cells from low-quality datasets (≤1 criterion), approximately 25% could not be reliably assigned to Martinotti (MC), non-Martinotti (nMC), or LRP subclasses (**Fig. 2i, 2j**), suggesting that insufficient integration can obscure underlying biological identities. In contrast, nearly all cells from datasets passing ≥2 criteria were confidently classified. Together, these results demonstrate that our three integrability criteria not only capture structural and batch-related consistency but also serve as stringent quality filters to exclude datasets with fewer transcriptomic features.

The parameters measured in Module 2 are generalizable, and can be applied to other datasets to assess integrability across different experimental conditions. We provide two examples. For the first example, we took unpublished snRNAseq datasets generated from 13 samples of mouse adult spinal cords, representing different experimental conditions from Aya Takeoka’s laboratory. One control sample, the one with the largest number of cells, was used as the “reference/base atlas”, akin to our Atlas-v0, and the integrability of subsequent biological samples were assessed using this reference sample. Even though cellular diversity, ages, and sample types were different, Module 2 can be used to assess integrability by computing the three criteria: mean anchor point (**Extended Data Fig. 3a, 3c),** fraction within k-NN (**Extended Data Fig. 3a, 3b, 3d)**, and k-BET rejection rate (**Extended Data Fig. 3a, 3e).**

**Figure 3.**
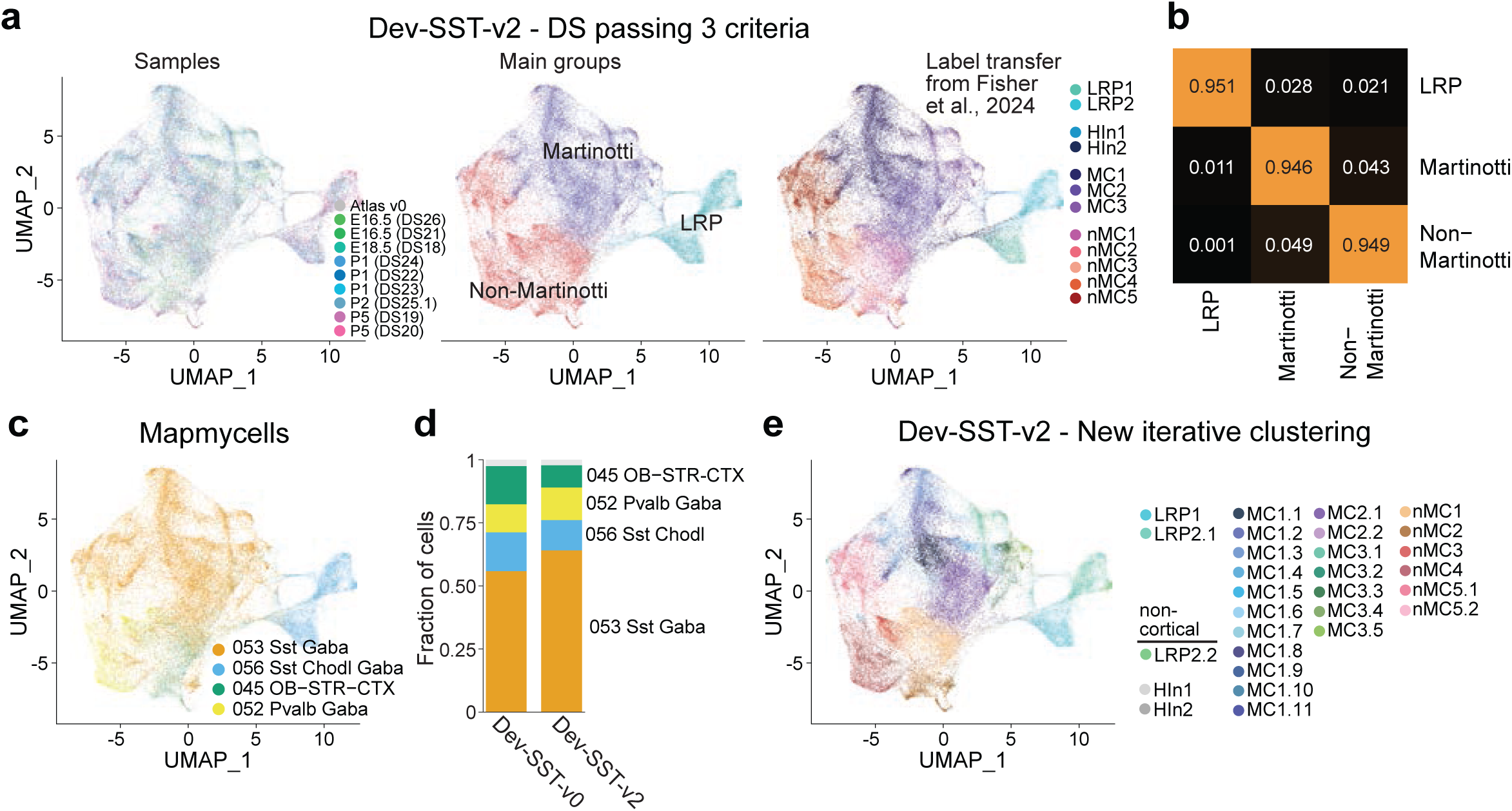
Dev-SST-v2 reveals 26 clusters of cortical SST+ inhibitory neurons. (**a**) UMAP visualization of Dev-SST-v2, showing 59,658 SST+ cells grouped into three main populations: Martinotti cells (MC), non-Martinotti cells (nMC), and long-range projecting neurons (LRP), across developmental stages (E16.5, E18.5, P1, P2, and P5). **(b**) Confusion matrix displaying high classification accuracy (>0.9 probability) for distinguishing MC, nMC, and LRP populations. (**c**) Fraction of cells in Dev-SST-v3 matched with high confidence (>0.9 probability) to adult cortical SST+ inhibitory neuron subclasses using the MapMyCells algorithm, with subclass annotations 053 Sst and 056 Sst-Chodl dominating. (d) Stacked bar plot comparing relative abundances of main cell types between Dev-SST-v0 and Dev-SST-v2, highlighting the stability of population structure across atlases. (**e**) Iterative clustering of Dev-SST-v2 reveals 26 stable clusters of cortical SST+ interneurons (>0.75 Jaccard index). Clustering results demonstrate the expansion of developmental diversity, identifying 18 MC types, 6 nMC types, and 2 LRP types (LRP1 and LRP2.1). Three rare clusters, HIn1, HIn2, and LRP2.2 clusters (<1% of all cells) are non-cortical.

For the second example, we constructed an atlas of developmental cortical PV+ interneurons, also a minority cell population. We examined 6 publicly available datasets (DS7.1, DS12.1-12.3, DS25.1-25.2), and determined the number of PV+ interneurons and used the dataset with the most numerous cell numbers as a base atlas (**Extended Data Fig. 4a, 4b**). Following the same pipeline, we measure anchor point, fraction of cells within k-NN (k=60), and k-BET test (**Extended Data Fig. 4b**). Then, we integrate 4 datasets that passed all 3 criterias (**Extended Data Fig. 4c**), and cluster based on iterative clustering to generate the atlas Dev-PV-v1 (**Extended Data Fig. 4d).** Using *Mapmycells*, we found that >85% of cells in Dev-PV-v1 are mapped to the adult 052 Pvalb Gaba with a probability of >0.9 (**Extended Data Fig. 4e, 4f)**. These two examples demonstrate that our criteria and pipeline are robust and generalizable.

### Integrability criteria do not depend on sequencing depth

To identify factors that influence integrability of independent datasets, we searched for dataset-specific features in experimental setup and computational analyses. Since all datasets were subjected to the same QC parameters in Module 1 (read counts, ribosomal RNA, mitochondria genes), cellular heterogeneity and general quality of transcripts within each dataset are unlikely to be the separating factors. We noted that sequencing read depth differs greatly between laboratories and samples, and thus examined whether this could be a potential factor.

To this end, we took two datasets that we generated (DS21 and DS24) from two stages, E16.5 and P1, and performed both shallow and deep sequencing on the same samples. For shallow sequencing, we targeted a sequencing depth of 5000 reads per cell; the actual mean reads achieved was 8,227 +/-794. For deep sequencing, we targeted 50,000 reads per cell; the actual mean reads achieved was 61,384 +/-20,484 (**Extended Data Fig. 5a**). Then, we performed the integrability test (Module 2) by quantifying the 3 parameters: anchor points with Atlas-v0, fraction of cells within k-NN, and the k-BET rejection test (**Extended Data Fig. 5b-5e)**. Despite approximately a 7.5 fold difference in sequencing depth between shallow and deep sequenced samples, both datasets passed the integrability criteria (**Extended Data Fig. 5d, 5e**). Our analysis thus indicates that sequencing depth alone is not enough to explain failed integration, and suggests that other experimental parameters, including sample preparation, may influence integrability.

### Dev-SST-v2 reveals 26 clusters of cortical SST+ inhibitory neurons

We integrated SST+ cells from 7 datasets that passed all 2 or 3 criteria (Dev-SST-v1) or only 3 criteria (Dev-SST-v2) of Module 2 into Atlas-v0 using the Seurat CCA function, and generated two atlases (**Extended Data Fig. 6a, 6b)**. Additional quality control steps were performed to remove doublet cells, stress cells, and clusters that have cells predominantly (>60% composition) from a single dataset (**Extended Data Fig. 1c** and see methods). After these exclusions, the final Dev-SST-v1 and Dev-SST-v2 have 63,572 (stages E16.5, E18.5, P1, P2, P5, and P7, P10) and 59,658 (stages E16.5, E18.5, P1, P2, and P5) SST+ cells respectively (**Extended Data Fig. 6a, 6b).**

To compare Dev-SST-v1 and Dev-SST-v2, we generated confusion matrices from the random forest predictions, summarizing how cells of each true type were assigned to predicted classes (**Extended Data Fig. 6c)**. The diagonal of each matrix represents correctly classified cells (diagonal, accuracy), whereas off-diagonal entries capture misclassifications (cross-talk) between related types. We then computed per-class differences (Δ) between the two atlas versions, v2-v1, for both diagonal accuracy and cross-talk to quantify relative changes in classification performance. Dev-SST-v2 showed consistently higher or maintained diagonal accuracies (**Extended Data Fig. 6d)** and reduced cross-talk (**Extended Data Fig. 6e)** across Martinotti and Non-Martinotti populations, supporting its use for subsequent analyses.

To assess whether we have sufficient sampling at each time point, we performed saturation analyses by subsampling from 5% to 95% (2,560 to 48,656 cells). At each step, we evaluated: (i) the fraction of conserved clusters, (ii) the fraction of conserved differentially expressed genes; (iii) the absolute difference in mean expression of all genes, as a measure of variation, and (iv) the robust Hausdorff distance which is quantifying the similarity between subsampled and full datasets (**Extended Data Fig. 6f**). We identified saturation points between 8,639 and 11,151 cells, based on the second derivative inflection points of these measures (**Extended Data Fig. 6f**). At the three main developmental grouped stages, E16.5, E18.5/P1, and P5, we have exceeded this threshold.

Dev-SST-v2 can be divided into three main cell populations—MC, nMC, and LRP (**Fig. 3a**)—in agreement with previous findings ^23^. These 3 populations can be discerned with high accuracy, ranging from 90-95% (**Fig. 3b**). Using *MapMyCells*, we found that, similar to Atlas-v0, most cells (> 75%) are matched with high confidence (>0.9 probability) to adult cortical SST+ inhibitory neurons with the subclass level annotation of 053 Sst, 056 Sst-Chodl (**Fig. 3c, 3d**). New iterative clustering of Dev-SST-v2 revealed 26 clusters (**Fig. 3e**) of stable cortical SST+ neurons, Jaccard index >0.75 **(Extended Data Fig. 6g)**, with about 3 times more clusters (**Fig. 3a**) than previously reported in these developmental stages ^23^. Each cluster has its cluster-specific gene expression profile (**Extended Data Fig. 7a, 7b, 7e)**.

Compared to our previous work ^23^, Dev-SST-v2 shows significantly more clusters of MCs (from 3 to 18) and nMCs (from 5 to 6) (**Fig. 3e)**. However, the number of LRP clusters stayed the same. We detected two types of cortical LRPs: LRP1 and LRP2.1, matching our previous work (**Fig. 3e**). LRP2.2 represents less than 3% of LRPs (**Extended Data Fig. 7c)**, and is a striatal interneuron contamination because of the strong expression of Nkx2-1 (**Extended Data Fig. 7c**), a transcription factor known to be expressed in early postnatal striatal SST+ interneurons but not cortical interneurons ^53^. The identification and functional validation of LRP2.1 highlight the pipeline’s utility in detecting rare, transient populations.

To validate if our CCA-based data integration of clusters from early stages (E16.5) correspond to those of later stages (E18.5/P1 and P5), we applied the Waddington Optimal Transport (WOT) framework, which explicitly models probabilistic transitions between timepoints (**Extended Data Fig. 8)**. The WOT analysis independently confirmed the inferred relationships: cells from E16.5 clusters showed the highest transition probabilities toward corresponding P1 clusters, and P1 cells toward P5 counterparts (**Extended Data Fig. 8a)**. The enrichment of strong diagonal couplings relative to off-diagonal ones (**Extended Data Fig. 8b -8d**) indicates that the observed developmental mappings are biologically robust rather than driven by integration artifacts.

### Cortical SST+ interneurons display heterogeneous modes of diversification

To determine when different types of SST+ interneurons emerge during development, we used *MapMyCells* to assign developmental MCs and nMCs to the adult subclasses and supertypes found in Yao et al. ^15^ (**Fig. 3a, 3c, Extended Data Fig. 9a - 9c**). Our premise is that an immature, developing SST+ neuron, sharing a similar transcriptomic signature than an adult SST+ cell, will likely become that subclass/supertype/type in adulthood. We set a high threshold for mapping, only considering matched transcriptomic pairs when the mapping probability is >0.9 (see methods).

At the subclass level, almost all (>99%) developing MCs are assigned to the adult subclass 053 Sst (**Extended Data Fig. 9a**). For nMCs, 85% of the cells mapped to their adult counterparts (**Extended Data Fig. 9a**). At the supertype level, the highest fraction of developing cells mapping to adult ones are found at P5, with 74% for both MC and nMC (**Extended Data Fig. 9b, 9c**). As early as P1, we were able to assign all 19 adult SST+ interneurons supertypes (0214 Sst Gaba_1 to 0232 Sst Gaba_19) to our developmental MC and nMC cells (**Extended Data Fig. 9d**).

Almost all the cells that were mapped to the 052 subclass Pvalb interneurons (**Fig 3c**) are nMC clusters, and they represent 26.5% of all nMCs (**Extended Data Fig. 9a**). The mapping of nMCs to the 052 Pvalb subclass may reflect the SST+ supertypes that are closely related to the transcriptomic and firing properties to Pvalb+ cells ^17,19^. In line with this, a recent preprint by the Allen Institute showed that during development, the Pvalb type 735, segregates from a branch of cells belonging to the Sst subclass ^54^. These findings, along with our analysis, suggest that some developmental SST+/nMCs clusters share more transcriptomic similarity with the adult Pvalb than the Sst supertypes.

Next, we examined whether the developmental SST+ interneuron clusters exhibit a one-to-one match to adult supertypes. We define a one-to-one match if >65% of cells in a developmental cluster map to a single adult supertype (probability of >0.9). Among the 18 developmental MC clusters, 7 have a one-to-one correspondence to adult Sst supertypes (**Fig. 4a, 4b**). or nMCs, in contrast, only 1 out of the 6 developmental clusters have a clear 1:1 match. Most developmental nMC clusters are mapped to multiple adult Sst supertypes (**Fig. 4c, 4d**).

**Figure 4.**
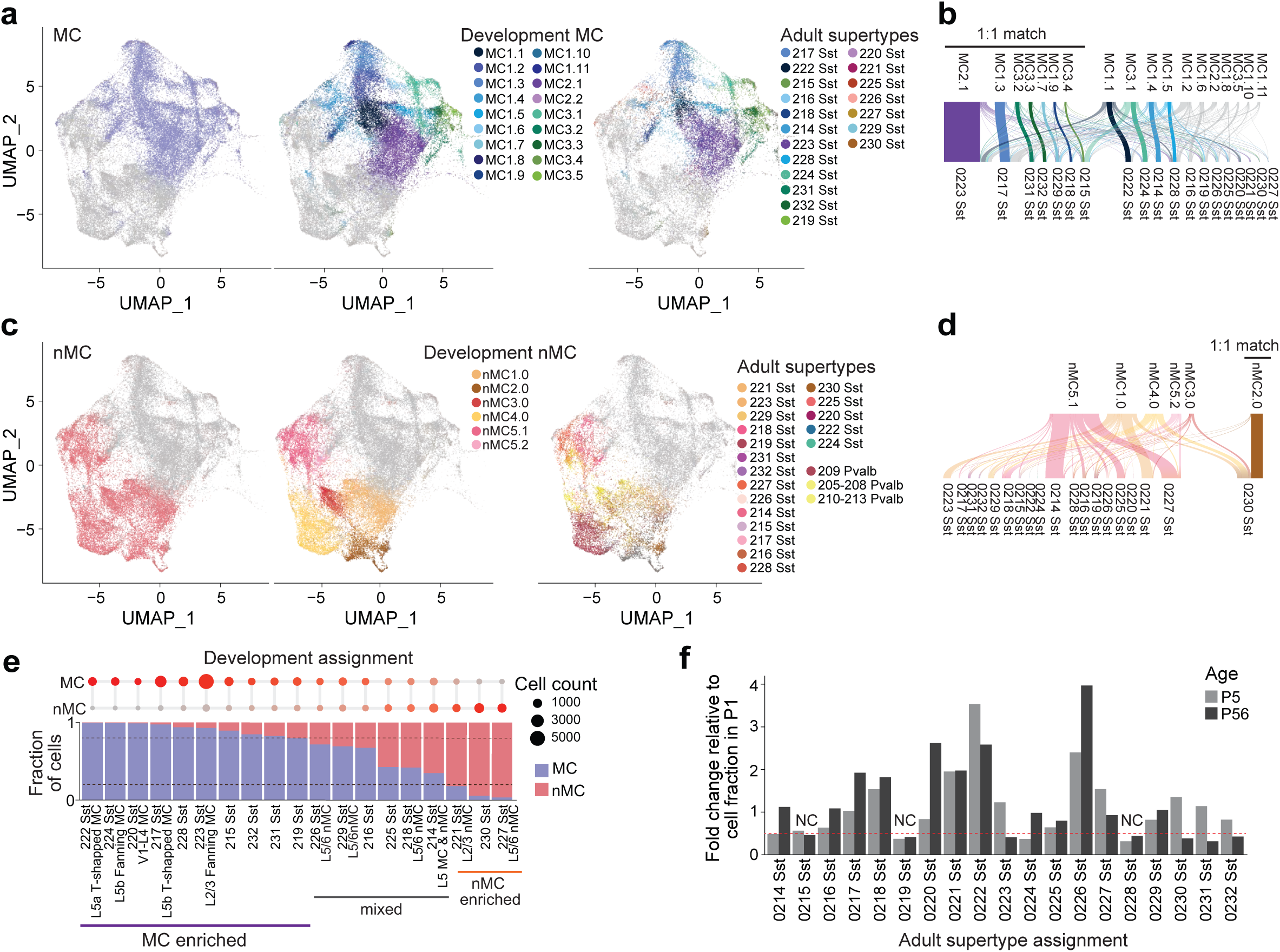
MCs and nMCs diversification during development. (a) UMAP of Dev-SST-v2, focusing on developmental interneuron clusters by adjusting the axis limits [UMAP 2: -5 to 5 instead of -5 to 10 in Fig 3]. Labeling highlights only MC for visualization. Developing MC clusters mapped to adult SST+ interneuron supertypes using MapMyCells with a high-confidence threshold (>0.9). Clusters with a one-to-one (1:1) match to adult supertypes are highlighted. (**b**) River plot of developmental MC clusters and their corresponding adult supertypes, illustrating 1:1 matches. (**c**) UMAP of developmental interneuron clusters by adjusting the axis limits [y-axis UMAP 2: -5 to 5 instead of -5 to 10 in Fig. 3]. Labeling highlights only nMC for visualization. nMC clusters mapped to adult SST+ supertypes, highlighting limited 1:1 matches. Notably, nMC clusters mapped to both SST+ and Pvalb+ adult subclasses. (**d**) River plot of developmental nMC clusters and their corresponding adult supertypes, illustrating a more flexible developmental trajectory with multiple clusters contributing to individual adult supertypes. Only 1 out of 6 nMC clusters showed a 1:1 relationship with adult supertypes. (**e**) Bubble chart (*top*) and bar plot showing the composition of adult SST+ supertypes derived from developmental MC and nMC clusters. Ten supertypes were exclusively composed of MC-derived cells, while three were exclusively nMC-derived. Supertypes with mixed compositions were primarily assigned to nMC adult types. (**f**) Fold changes in the relative abundance of each Sst supertype from P1 to P5 and P1 to P56.

For each of the Sst supertypes assigned to developmental interneurons, we also calculated the relative fraction composed of either MC or nMC clusters (**Fig. 4e**). We found that 10 out of the 19 supertypes were exclusively composed of MCs (relative fraction >75%), while only 3 out of the 19 supertypes consisted of nMCs (**Fig. 4e**). The Sst supertypes that are exclusively composed of developmental MCs are also well defined as MCs in the adult, based on morphology, electrophysiology, transcriptomics, (MET types) and spatial distributions ^19^ (**Fig 4e**).

For example, supertype 222 Sst, MET-7 in Gouwens et al. ^19^, is a L5a T-shaped MC; 224 Sst - MET-5, is a L5 fanning shaped MC, 220 Sst, MET-8 is a L4 MC; 217 Sst, MET-6, is L5b T-shaped MC; and 223 Sst, MET-3 is a L2/3 fanning MC ^16,19,54^. For the 3/19 supertypes that are exclusively nMC, these include supertypes 221 and 227 which represent MET-2 L2/3 nMC with fast-spiking properties and MET-12/13, L5/6 nMC respectively (**Fig. 4e**). In contrast, the supertypes that show mixed developmental MC and nMC compositions are mostly assigned to nMC adult types. These analyses show that while most adult MCs emerge from developmental MC clusters, the diversification of adult nMCs may be more flexible, consisting of both developmental MC and nMC clusters.

To assess the biological relevance of the identified clusters, we performed MERFISH on P5 tissue using a customized probe set comprising approximately 500 genes (**Extended Data Table 2**), designed to classify our developmental clusters with high accuracy (**Extended Data Fig. 10a**). Most clusters displayed distinct spatial distributions within the cortex (**Extended Data Fig. 10b**). Six clusters were confined to infragranular cortical layers (**Extended Data Fig. 10c**), while five were restricted to ventral regions such as the insular and piriform cortices (**Extended Data Fig. 10d**).

As we observed that several developmental clusters corresponded one-to-one with adult Sst supertypes (**Fig. 4**), we next asked whether their spatial localization at P5 resembled that observed in the adult. Using adult MERFISH data from Zhang et al. ^20^, and our P5 data, we compared the spatial distribution of eight developmental clusters: four from infragranular cortical layers and four from the ventral cortex. All eight developmental clusters and their corresponding adult supertypes exhibited similar spatial localization (**Extended Data Fig. 10e,f)**. The pronounced anatomical specificity of these developmental clusters, identified by our Dev-SST-v2 analysis, indicates that dorsal and ventral cortical SST⁺ populations represent distinct cell types. Moreover, the spatial segregation of ventral interneurons (**Extended Data Fig. 10f**) by P5, prior to the onset of programmed cell death, suggests that this organization emerges during migratory processes. Collectively, these results demonstrate that the granularity of our clustering approach captures biologically meaningful anatomical and regional distinctions.

We then sought to determine when the relative abundance of SST+ interneurons is established during development. To address this, we analyzed the relative occurrence of each SST+ supertype at different developmental stages. We first quantified the relative fraction of each Sst supertype in the adult cortex using published data from Yao et al. (P56) ^15^. We then assessed the relative abundance of each Sst supertype at earlier developmental stages (P1 and P5) within both major classes: MC and nMC. To track developmental changes, we calculated the fold change in relative abundance for each Sst supertype at P5 and P56 relative to P1 (**Fig. 4f**). Our analysis revealed that, with the exception of three supertypes (0215, 0219, and 0228; **Fig. 4f**), most Sst supertypes undergo substantial shifts in relative abundance between P1 and P5, as well as between P1 and P56 (>0.5 fold change). This suggests that the relative proportions of most SST+ supertypes are not fixed early in development but instead continue to change throughout postnatal maturation. This finding is consistent with observations from a recent preprint ^54^. Overall, our data indicate that some Sst supertypes emerge relatively late in development, through several distinct diversification trajectories.

### LRP1 and LRP2.1 are distinct cell types with divergent trajectories

Our previous study ^23^, using MetaNeighbor analysis ^55^, linked developmental LRP1 to the adult Sst-Chodl-64 and Sst-Chodl-65 subtypes, while LRP2.1 (formerly LRP2) was associated with Sst-Chodl-63 in the Yao et al. (2021) ^17^ datasets. To refine this classification to the new ABC Atlas Yao et al. (2023) ^15^, we again used *MapMyCells* for single-cell mapping. This analysis revealed that LRP1 corresponds to the Sst-Chodl-Gaba-2 (supertype 241), whereas LRP2.1 aligns with Sst-Chodl-Gaba-4 (supertype 239) (**Fig. 5a**).

**Figure 5.**
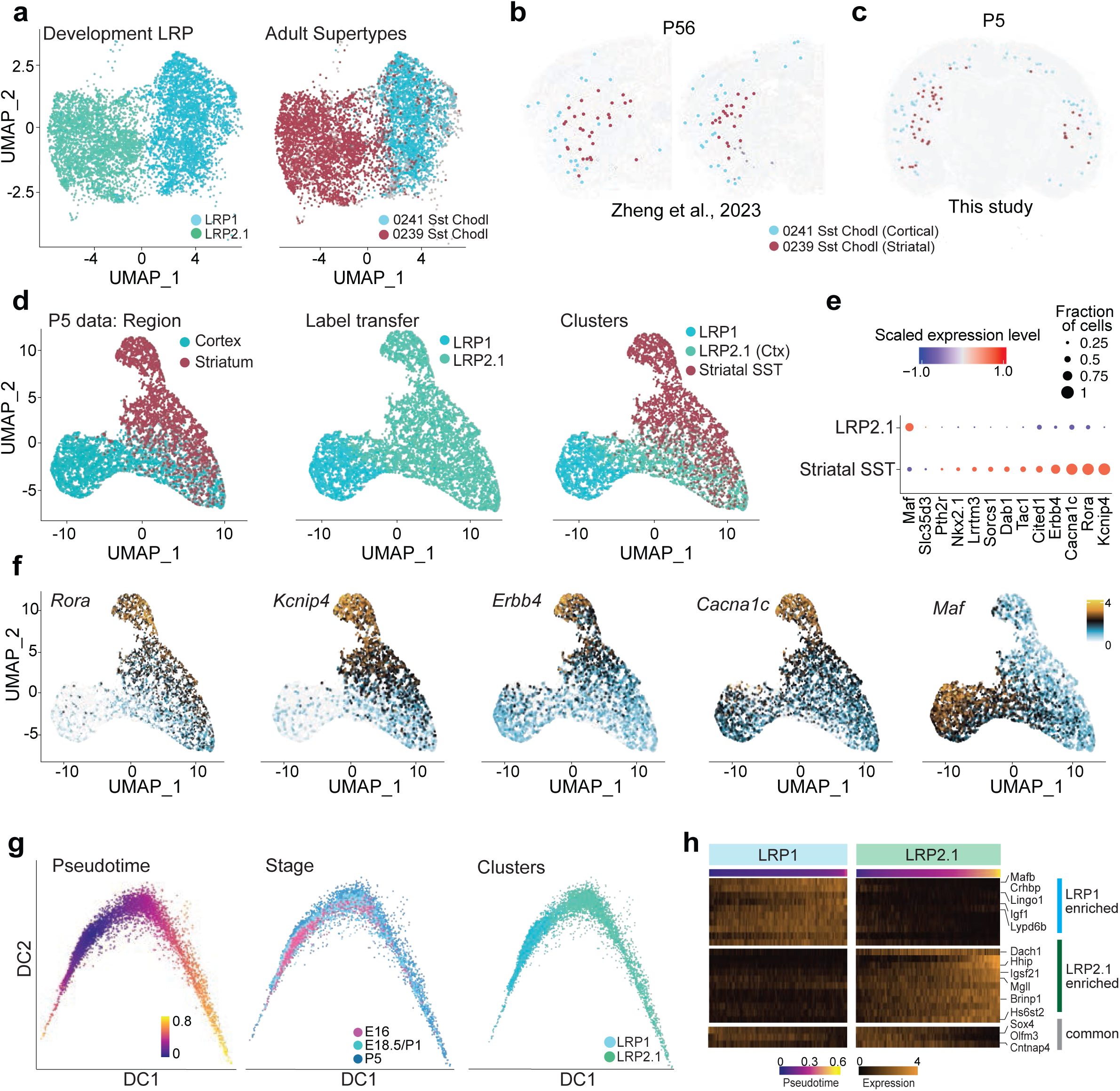
LRP1 and LRP2.1 are distinct cortical cell types. (**a**) UMAP of developmental LRP clusters, highlighting distinct mappings of LRP1 to adult supertype 241 Sst-Chodl-Gaba-2 and LRP2.1 to adult supertype 239 Sst-Chodl-Gaba-4. (**b**) Adult spatial distribution of LRP supertypes in the ABC Atlas MERFISH data (Zhang et al., 2023)^20^. Note that Supertype 241 Sst-Chodl-Gaba-2 is exclusively cortical, while supertype 239 Sst-Chodl-Gaba-4 is restricted to the striatum. (**c**) P5 Spatial distribution of P5 LRP supertypes using customized MERFISH data from this study showing mapping of 0239 Sst-Chodl-Gaba-4 in the cortex. (**d**) UMAP of P5 cortical LRP clusters and striatal SST+ interneurons labeled by region of tissue, label transfer from Dev-SST-v2, and cell cluster. (**e**) Bubble plot of key marker genes distinguishing LRP2.1 and striatal SST+ interneurons. (**f**) UMAP showing expression of key marker genes in cortical LPRs and striatal SST+ interneurons. (**g**) Pseudotime analysis of LRP1 and LRP2.1 using diffusion map space and developmental stage data, showing two distinct branches. (**h**) Expression of shared and branch-specific genes along pseudotime progression.

Further spatial mapping with the ABC Atlas MERFISH ^20^ showed a striking anatomical segregation: Sst-Chodl-Gaba-2 (supertype 241) is confined to the cortex, while Sst-Chodl-Gaba-4 (supertype 239) is found exclusively in the striatum (**Fig. 5b**). In contrast, at P5, MERFISH data from this study reveal that Sst-Chodl-Gaba-4 (supertype 239) can be found in the cortex (**Fig. 5c**). Notably, Sst-Chodl-63 from the Yao et al. (2021) dataset ^17^, which accounts for less than 5% of all LRPs, also maps to Sst-Chodl-Gaba-4, further indicating that LRP2.1 cells possess a transcriptomic signature characteristic of adult striatal interneurons (**Extended Data Fig. 11a**; see Discussion).

Given that LPR2.1 cells map transcriptomically to adult striatal interneurons, we next assess whether they are transcriptomically different clusters from striatal SST+/Nos1+ interneurons. To test this, we isolated SST+ striatal interneurons at P5 (**Fig. 5d**) and performed single-cell RNA sequencing. We integrated these data with all P5 LPR cells annotated in the Dev-SST-v2 dataset (**Fig. 5d**), and generated a joint UMAP embedding. We found that cortical LPR2.1 cells clustered in close proximity to both cortical LPR1 cells—previously defined long-range GABAergic neurons—and striatal SST+ interneurons (**Fig. 5d**). Although SST+ striatal interneurons shared several transcriptomic features with cortical LPR2.1 cells, including being assigned an LPR2.1 by label transfer approaches using the labels from Dev-SST-v2 (**Fig 5d**, middle panel), the two populations exhibited distinct cluster boundaries (**Fig 5d**, right panel) and differential expression of key genes such as *Rora*, *Kncip4*, *Erbb4*, *Cacna1c*, and *Maf* (**Fig. 5e, 5f**). Together, these analyses indicate that LPR2.1 have partial transcriptomic overlap with cortical LPR1 and striatal SST+ interneurons. However, LPR2.1 is molecularly distinct from these two cell clusters.

Having established that cortical LPR2.1 cells constitute a distinct cortical population, we next asked whether they follow a developmental trajectory separate from that of LPR1 cells. To determine whether LRP1 and LRP2.1 follow distinct developmental trajectories, we performed pseudotime analysis using the expanded Dev-SST-v2 dataset. First, we identified key genes required for constructing the pseudotime trajectory. Using the ANTLER algorithm ^56^, we identified 335 genes grouped into four distinct modules involved in neuronal differentiation, axon development, and synapse assembly (**Extended Data Table 3**; see Methods). We then constructed the pseudotime trajectory by generating diffusion maps, which revealed two clearly separated branches corresponding to LRP1 and LRP2.1 (**Fig. 5g**). Even at early pseudotime points, these two clusters were distinct, each expressing a unique set of branch-specific genes (**Fig. 5h**). Taken together, our findings indicate that LRP1 and LRP2.1 follow distinct developmental trajectories and map to different adult Sst-Chodl supertypes. These results suggest that LRP1 and LRP2.1 represent two fundamentally distinct cortical cell types rather than different cell states.

### LRP2.1 Is a transient cell population that declines between P5 and adulthood in the mouse cortex

Our analyses indicate that most LRP2.1 cells do not persist in the adult neocortex, despite their unique developmental trajectory. This is supported by the adult cortical transcriptomic dataset from Yao et al. (2021) ^17^ (**Extended Data Fig. 11a**) and spatial transcriptomic atlas data ^20^ (**Fig. 5b, 5c**), both of which show that fewer than 5% of LRPs map to LRP2.1 in the adult brain (**Extended Data Fig. 11a**). In contrast, the Dev-SST-v2 reveals that LRP2.1 constitutes approximately 25% of all LRPs at E16.5, increasing to ∼55% by P5 (**Fig. 6a**). Given that cortical inhibitory neurons do not migrate back to the subpallium or striatum after reaching the cortex ^9,57–59^, we hypothesized that LRP2.1 is a transient cell type that is lost by adulthood.

**Figure 6.**
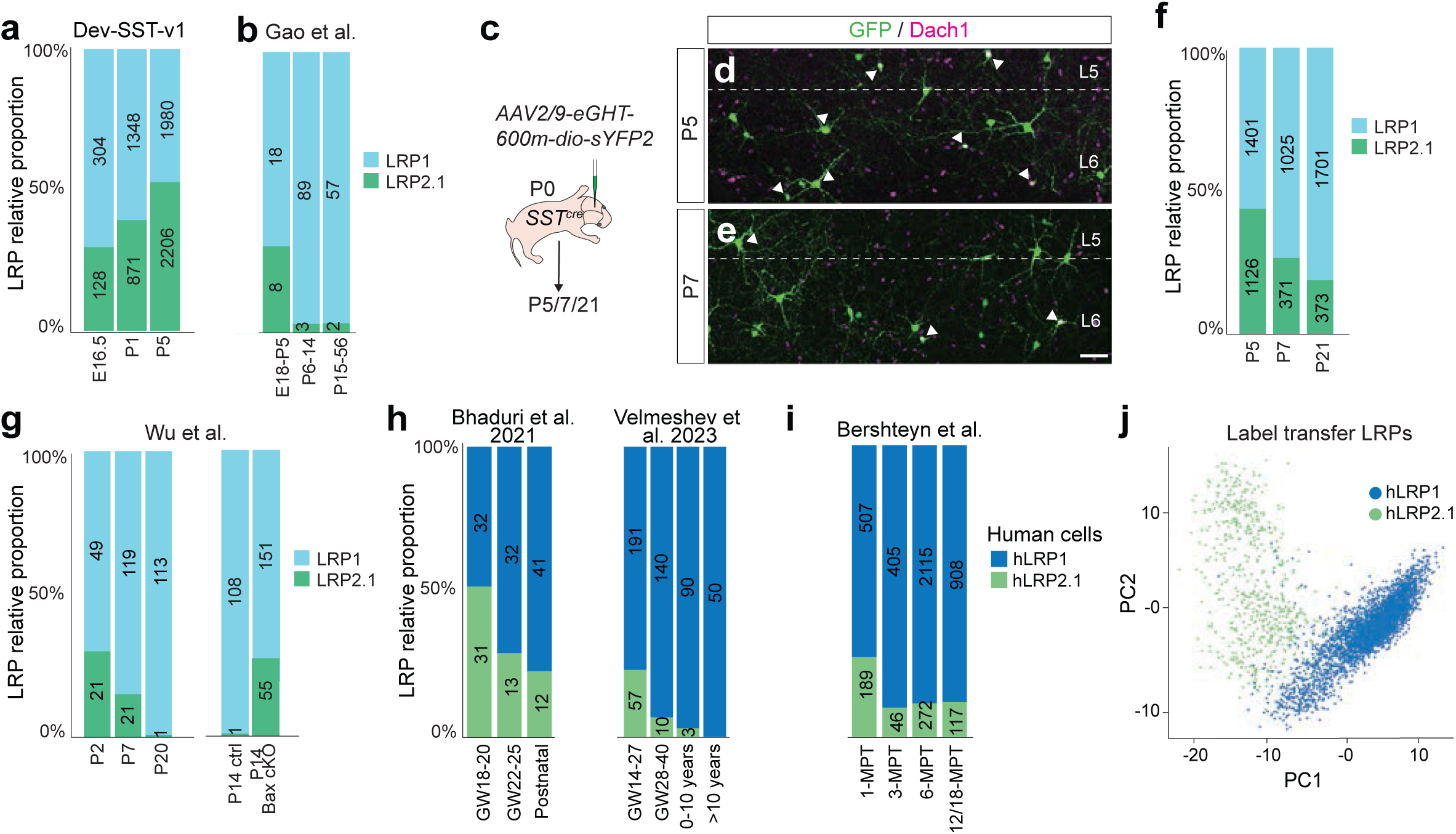
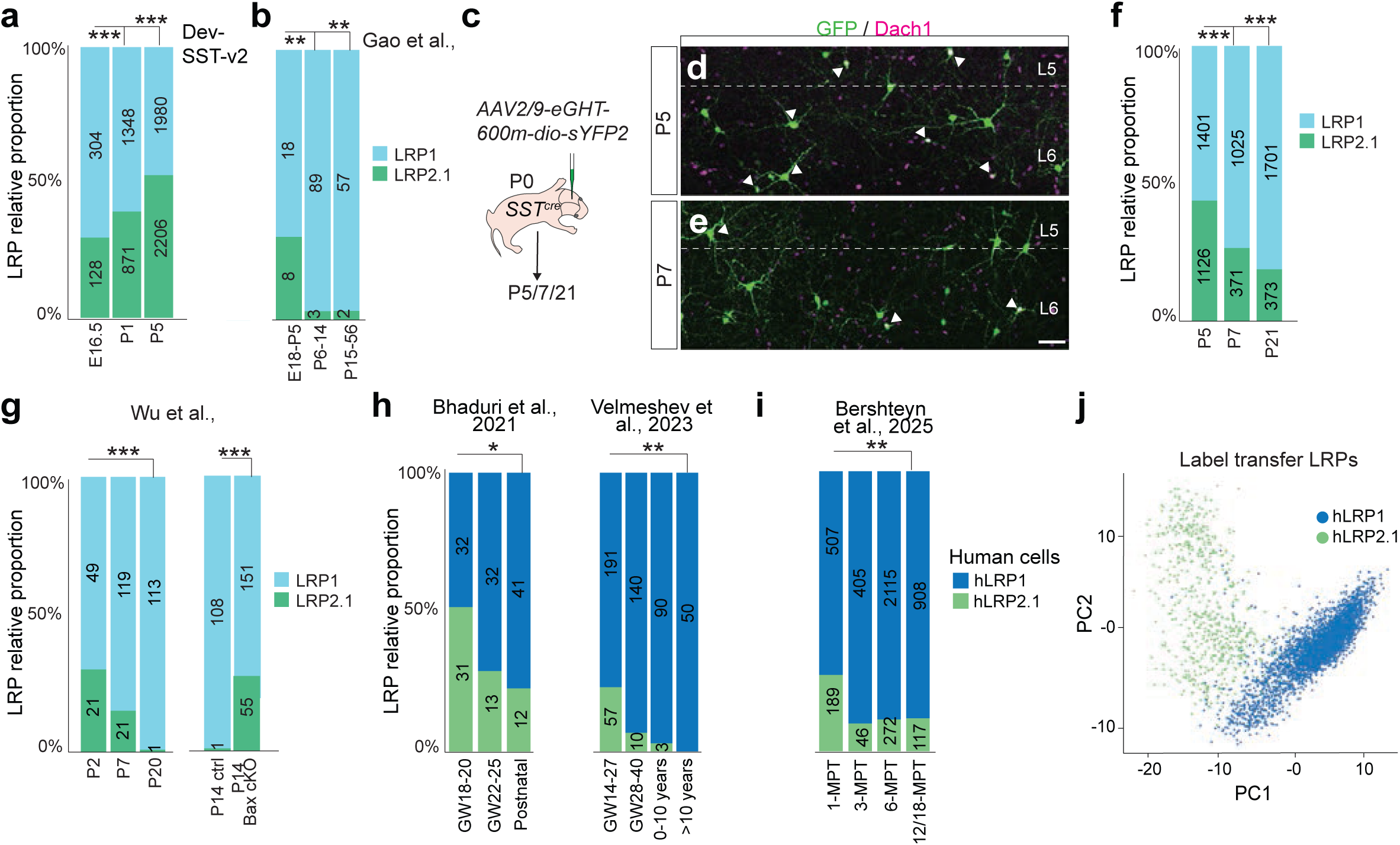
LRP2.1 is a transient cell type in both mouse and human neocortex. **(a**) Relative proportions of LRP1 and LRP2.1 in Dev-SST-v2. (**b**) Validation of LRP1 and LRP2.1 relative abundance, reanalysed from Gao et al. 2024, bioRxiv ^54^. (**c**) Experimental schema of labeling LRPs by AAV2/9-eGHT-600m-dio-sYPF2 in SSTcre mice at P0. (**d**) Co-immunostaining for GFP and Dach1 (LRP2.1 marker) at P5 and (**e**) P7. White arrow heads indicate double positive cells (LRP2.1). Scale bar =100µm. (**f**) Proportions of LRP1 and LRP2.1 quantified by immunostaining at P5, P7 and P21, n = 3 animals per time point. (**g**) Relative abundance of LRP1 and LRP2.1 at P2, P7, and P14 reanalysed from control mice and P14 in Bax cKO mice, datasets from Wu et al., 2024 bioRxiv ^68^. (**h**) Relative abundance of human LRP1 and LRP2.1 in developing neocortex reanalysed from independent datasets (Bhaduri et al ^69^., and Velmeshev et al. ^70^) (**i**) Relative abundance of LRP1 and LRP2.1 in hPSCs-derived MGE interneuron xenotransplanted in the mouse neocortex, reanalysed from Bershteyn et al.^71^ The number on each bar represents the number of LRPs found in each stage. (**j**) PCA plot of hLRP1 and hLRP2.1, reanalysed from Bershteyn et al.^71^ * p<0.05, ** p < 0.01, *** p< 0.001.

To track these populations beyond P5, we performed label transfer analysis on snRNA-seq datasets from a recent preprint by the Allen Institute (Gao et al.) ^54^, which spans developmental stages E13.5 to P60. Although LRPs were relatively rare in these datasets, we observed that LRP2.1 abundance peaks at P5 before rapidly declining, stabilizing to less than 10% in adulthood (**Fig. 6b**). To ensure that this sharp reduction at P7 does not reflect selective vulnerability of these cells during tissue dissociation, we examined LRP1 and LRP2.1 abundance in the cortex at P5, P7, and P21 using histology. We labeled LRPs with a modified enhancer virus targeting the *Nos1* locus, AAV2/9-eHGT-600m-sYPF2 ^60^. In early postnatal stages, we noted that this enhancer also transiently labels pyramidal neurons due to the low and transient expression of Nos1 during development ^61^ (**Extended Data Fig. 11b**). Therefore, we restricted expression to SST+/NOS1+ neurons by subcloning the enhancer into a Cre-dependent construct, and generated pAAV-eHGT-600m-dio-sYPF2 (**Extended Data Fig. 11c**). We injected AAV2/9-eHGT-600m-dio-sYPF2 into the lateral ventricles of P0 *Sst^Cre^* mice and performed immunostaining at P5, P7, and P21 (**Fig. 6c**), and confirmed that all fluorescently labeled cells (sYPF2+) were NOS1+ (**Extended Data Fig. 11c**).

Further data mining from the Dev-SST-v2 atlas revealed that a combination of three markers—either *Sst/Dach1/Nos1* or *Sst/Mafb/Nos1*—distinguishes LRP1 and LRP2.1 with >80% accuracy (**Extended Data Figs. 11d, 11h**). To differentiate between LRP1 and LRP2.1 in histological sections, we performed co-immunostaining with GFP/Dach1, a positive marker for LRP2.1 (**Extended Data Figs. 11e, 11i**), and GFP/MafB, a positive marker for LRP1 (**Extended Data Figs. 11f, 11j**). Control experiments confirmed that GFP/Dach1 and GFP/MafB staining did not significantly alter LRP1 and LRP2.1 quantification (**Extended Data Figs. 11g, 11k**).

Immunostaining revealed that at P5, LRP1 and LRP2.1 were present in nearly equal proportions, consistent with Dev-SST-v2 quantifications (**Fig. 6a, 6d, 6e, 6f**). By P7, however, LRP2.1 abundance decreased substantially (**Fig. 6e, 6f**), and by P21, only ∼18% of LRPs were LRP2.1-positive (***p<0.001, **Fig. 6f**). This is consistent with our previous findings ^23^, which showed that only 10% of LRPs were Dach1+ in the motor, somatosensory, and visual cortex at P28. Together, these results demonstrate that LRP1 and LRP2.1 exist in nearly equal proportions at P5. However, by adulthood, LRP2.1 has largely disappeared, supporting the hypothesis that this cell population is transient and selectively eliminated after birth.

### Programmed Cell Death Selectively Eliminates LRP2.1 in the Mouse Cortex

In the mouse neocortex, postnatal programmed cell death removes approximately 40% of cortical GABAergic neurons ^62–65^. Traditionally, this process has been considered non-selective, affecting all interneuron subtypes uniformly ^62–64,66,67^. The rapid decline of LRP2.1 between P5 and P7 does not align with this general timeline, as most studies report that interneuron loss primarily occurs between P7 and P12 ^62–65^. This discrepancy raises the question of whether LRP2.1 undergoes selective elimination through a distinct developmental mechanism.

Recent preprint by the Fishell lab, Wu et al. ^68^, suggests that cell-type-specific postnatal programmed cell death may be possible. Their study on Fezf2-deficient mice, in which layer 5 excitatory pyramidal neurons are absent and layer 6 excitatory neurons are supernumerary—showed that some L5/6 SST+ interneuron subtypes adjust their relative abundance through selective programmed cell death. This finding challenges the traditional view that PCD is a uniform process and suggests that specific interneuron populations can be targeted for elimination in postnatal circuits. We therefore hypothesized that a similar mechanism shapes the composition and diversity of LRP populations.

To investigate whether programmed cell death selectively eliminates LRP2.1, we re-analyzed snRNA-seq datasets from Wu et al. ^68^, which included inhibitory neurons from control (*Nkx2.1^Cre^;Bax^fl/+^;Rosa26^LSL-h2b-GFP^*) mice at P2, P7, and P14, as well as Bax cKO (*Nkx2.1^Cre^;Bax^fl/fl^;Bak^-/-^;Rosa26^LSL-h2b-GFP^*) mice at P14, in which interneurons do not undergo programmed cell death. In wild-type controls, LRP2.1 abundance declines sharply between P2 and P7, disappearing entirely by P14 (**Fig. 6g**). In Bax cKO mice, however, LRP2.1 persists at P14 at levels comparable to P2 (∼25%), strongly indicating that its loss in wild-type animals is due to programmed cell death. Importantly, no significant changes were observed in the abundance of other SST+ interneuron supertypes (**Extended Data Fig. 11l**), further supporting the conclusion that LRP2.1 is selectively eliminated by programmed cell death. Together, these findings demonstrate that LRP2.1 is uniquely targeted for elimination via programmed cell death, providing the first evidence that inhibitory neuronal diversity can be sculpted through cell-type-specific postnatal elimination.

### LRP2.1 Emerges in the Human Developing Cortex and Declines by Birth

To determine whether LRP2.1 is conserved in humans and is eliminated during development, we re-analyzed scRNA-seq datasets from fetal, pediatric, and adult human cortex from two independent studies ^69,70^. Using label transfer on cell clusters expressing *Sst+/Nos1+/Npy+*, we identified human LRP1 (hLRP1) and LRP2.1 (hLRP2.1).

Similar to mice, hLRP1 and hLRP2.1 were present in nearly equal proportions at early stages. hLRP2.1 constituted ∼50% of LRPs at gestational weeks (GW) 18–20 but declined sharply by birth (**Fig. 6h**). Re-analysis of snRNA-seq data from hPSC-derived MGE interneurons xenotransplanted into mice ^71^ confirmed this trend: hLRP2.1 was most abundant at 1 month post-transplantation (1-MPT) but declined sharply by 3-MPT (**Fig. 6i, 6j, Extended Data Fig. 11m**).

Since programmed cell death in the human central nervous system begins around GW20 and concludes by birth ^72^, our findings strongly suggest that hLRP2.1 is selectively eliminated during this period. These findings indicate a conserved mechanism across species, highlighting selective removal of LRP2.1 by cell-type-specific programmed cell death.

### At P4–P6, LRP2.1 neurons are functionally connected to both cortical and thalamic excitatory inputs

To functionally validate the transient cell type LRP2.1, we investigated its excitatory presynaptic partners across postnatal development. Previous work has shown that layer 5 SST⁺ neurons receive transient thalamic input during early postnatal stages ^6,73^. In particular, Tuncdemir et al., ^6^ demonstrated that thalamic neurons form monosynaptic connections with SST⁺ interneurons when rabies pseudoviruses (RV) are injected during the first postnatal week. We therefore asked whether LRP neurons at P5 also receive thalamic input similar to SST⁺ interneurons

To test this, we injected *SST^cre^* mice at P0 with two types of AAV-helper viruses carrying Cre-dependent expression cassettes for GFP, optimized rabies G protein, and the envelope receptor, TVA. One helper virus used a general synapsin promoter (AAV2/9-syn-Flex-nGToG - nuclear GFP, TVA, and optimized G-protein) to label all pan SST+ cells (interneurons and LPRs), whereas the other employed an LPR-specific enhancer (AAV2/9-600m-Flex-nGToG) to selectively label LPRs. At P4, we injected an RV expressing TdTomato and Flp recombinase (N2c-ΔG-Tdt-Flp) into the somatosensory cortex and performed histological analyses of TdTomato+ presynaptic neurons, normalizing to the number of starter cells (TdTomato+/GFP+ cells - double positive) per brain at P8 (**Extended Data Fig. 12a**).

Consistent with previous findings, *SST^Cre^* mice injected with the synapsin-promoter helper virus (starter cells = all SST+ cortical cells) showed extensive labeling of thalamic and deep-layer cortical neurons (**Extended Data Fig. 12b, 12c**), replicating the result from Tuncdemir et al ^6^. Mice injected with the LPR-specific helper virus, while having less number of starter cells, exhibited extensive labeling of thalamic neurons, deep-layer cortical neurons, and subplate neurons (**Extended Data Fig. 12e, 12f)**. We compared the density presynaptic input cells (Tdt+ only per mm^2^ tissue) per 100 starter cells in the brains with either SST+ or LPRs as starters and found that there were significantly more cells in the subplate (** p<0.01) and thalamus (***p<0.001) when LRPs were starters (**Extended Data Fig. 12d, 12g)**. These results indicate that, similar to SST+ interneurons ^6^, LRPs receive substantial thalamic input and are monosynaptically connected to subplate neurons during early postnatal development.

We next compared thalamic innervation between LRP1 and LRP2.1 neurons. We labeled LRP neurons using an enhancer-driven AAV (AAV2/9-eHGT-600m-sYPF2) and performed co-immunostaining for GFP, Dach1 (a marker for LRP2.1), and VgluT2, which marks thalamic presynaptic terminals, at P4, P8, and P14 (**Fig. 7a – 7e**). At P4, both subtypes exhibited dense VgluT2⁺ puncta surrounding their soma and proximal dendrites, indicative of strong thalamic input (**Fig. 7a, 7c, 7e**). This thalamic innervation markedly declined by P8 and was largely absent by P14 (**Fig. 7b, 7d, 7e**), consistent with a transient thalamic connection previously reported for L5b SST+ cells ^6^.

**Figure 7.**
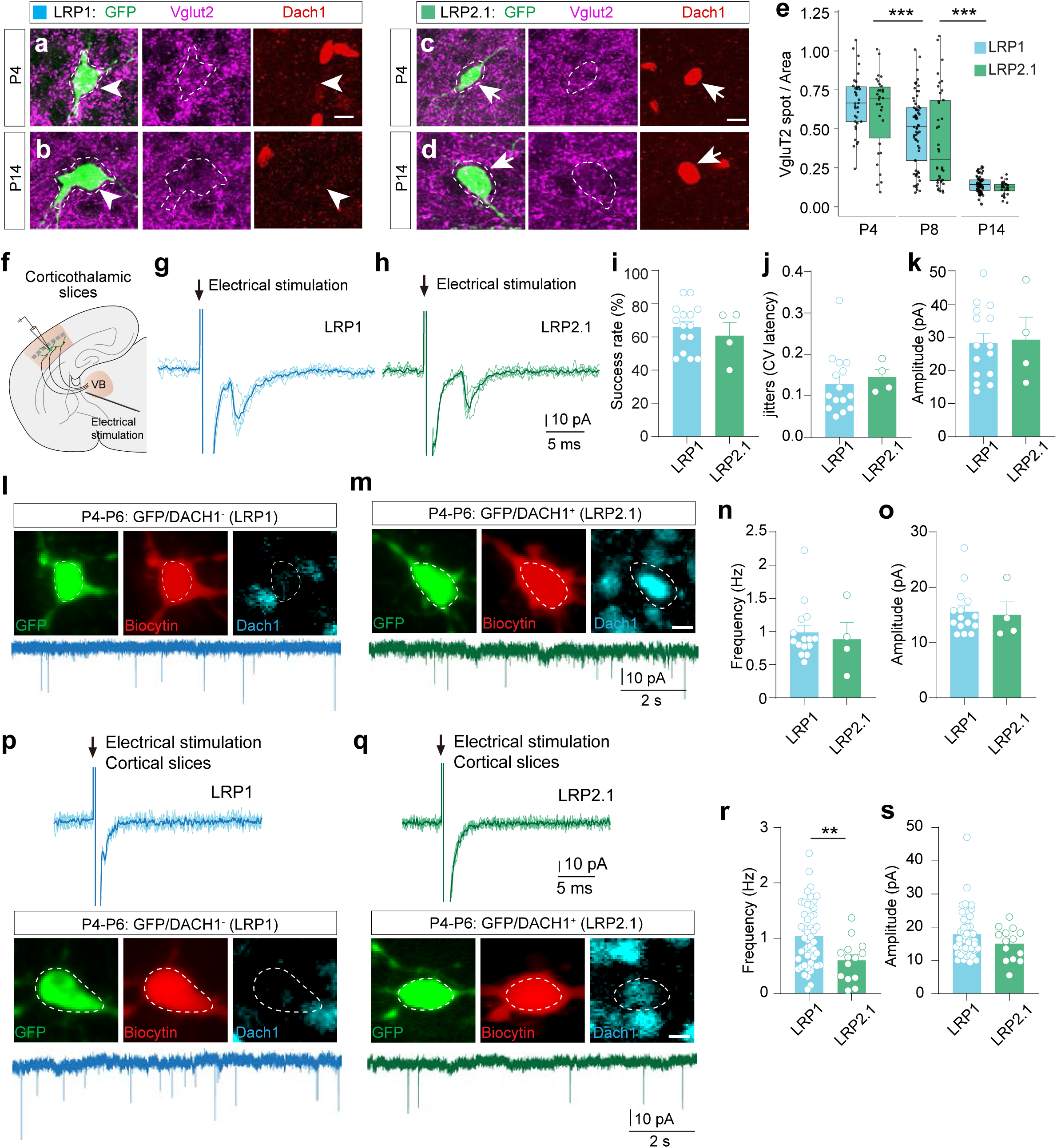
LRP1 and LRP2.1 neurons receive comparable transient thalamic input but differ in cortical excitatory connectivity. (a–d) Confocal images of LRP neurons labeled with an enhancer-driven AAV (AAV2/9-eHGT-600m-sYPF2) and immunostained for GFP, VgluT2, and Dach1 at P4 and P14. Both LRP1 (**a, b**: GFP⁺/Dach1⁻) and LRP2.1, n = 3 - 4 animals per conditions, >20 cells per brain (**c, d**: GFP⁺/Dach1⁺) neurons exhibited dense VgluT2⁺ perisomatic puncta at P4, which declined markedly by P8 and were largely absent by P14. (**e)** Quantification of VgluT2⁺ puncta density (spot number per um^2^ cell area) across ages, ***p<0.001 Two-factor ANOVA. (**f)** Schematic of thalamocortical slice preparation. (**g, h)** Representative thalamus-evoked EPSCs in LRP1 and LRP2.1 neurons at P5–P6. (**i–k)** Evoked EPSC success rate, latency jitter (coefficient of variance of latency), and evoked EPSC amplitude did not differ between subtypes. (**l, m)** Representative traces of spontaneous EPSCs (sEPSCs) recorded in thalamocortical slices, with post-hoc immunostaining confirming subtype identity. (**n, o)** sEPSC frequency and amplitude were similar between LRP1 and LRP2.1. (**p, q)** Cortical slices with severed thalamic afferents, thalamic stimulation failed to elicit EPSCs in either subtype. (**r)** sEPSC frequency from cortical slices with severed thalamic afferents was significantly lower in LRP2.1 neurons, **p<0.01. (**s**) sEPSC amplitude remained unchanged, indicating reduced cortical excitatory input to LRP2.1 cells. (**i–k, n, o, r, s)** Each dot is a single patched cell, from n > 3 animals per condition.

To assess the functional strength of these inputs, we performed whole-cell patch-clamp recordings from labeled LRP neurons in thalamocortical slices preserving corticothalamic connectivity (**Fig. 7f - 7h**). At P5-P6, electrical stimulation of the thalamus reliably evoked excitatory postsynaptic currents (EPSCs) in both LRP1 (GFP⁺/Dach1⁻) and LRP2.1 (GFP⁺/Dach1⁺) neurons with similar success rate (**Fig. 7i**), jitters (**Fig. 7j**, coefficient of variance of latency ^74^), and evoke EPSC amplitudes (**Fig. 7k**). The frequency and amplitude of spontaneous EPSCs (sEPSCs), which reflect combined thalamic and cortical excitatory inputs, did not differ between the two subtypes (Fig. **7l** **– 7o**).

Since our rabies tracing indicated that LRPs also received monosynaptic subplate input, we asked if LRP1 and LRP2.1 differ in their cortical excitatory inputs. For that, we performed patch-clamp experiments in cortical slices, in which we severed the thalamocortical afferents. We first verified the loss of thalamic connection by stimulating the thalamus, and failing to elicit EPSCs in either LRP subtypes (**Fig. 7p, 7q**). Under these conditions, the frequency of sEPSCs was significantly reduced in LRP2.1 neurons compared with LRP1 (**Fig. 7r**), while amplitude remained similar (**Fig. 7s**). These results suggest that although both LRP1 and LRP2.1 receive comparable thalamic input, LRP2.1 neurons may have fewer cortical excitatory connections. Together, these findings indicate that LRP2.1 neurons are transiently integrated into both cortical and thalamic excitatory circuits during early postnatal development.

## Discussion

### A modular computational pipeline for analyzing rare cell types

The challenge in studying rare cell types is their sparse representation within individual scRNA-seq experiments necessitating the aggregation of multiple independent datasets for proper sampling. However, integrating datasets with significant technical variability risks conflating biological variation with batch effects. Our bioinformatics suite addresses these two key issues by (1) isolating specific cell types or subclasses from diverse datasets (both published and unpublished) and (2) systematically evaluating whether datasets generated under different experimental conditions can be reliably integrated.

In Module 1, we performed a marker-informed enrichment rather than a fully automated classification. Populations are selected based on canonical markers (e.g., Sst, Lhx6), applied either at the single-cell or cluster level, which allows the inclusion of developing SST⁺ precursors whose Sst expression may be transient or low. This approach minimizes false exclusions that would occur with strict marker thresholds while maintaining specificity, as confirmed by subsequent MapMyCell analysis showing that >75% of selected cells map with >0.9 probability to SST⁺ subclasses (“053 SST” and “056 SST Chodl”).

Our integrability test (Module 2) quantifies structural similarities between datasets to ensure integration is performed only on datasets where batch effects are proportional to those in our reference atlas (Dev-SST-v0), thereby minimizing overcorrection while preserving biologically meaningful variation. Striking this balance is critical for rare cells, which are often clustered into a single group due to over-smoothing during batch correction—a significant issue in the field that has been highlighted by several methodological studies ^75–77^.

The iterative clustering procedure (Module 3) was performed independently of adult cell-type annotations, ensuring that developmental organization emerged directly from the data rather than from prior assumptions. This approach allowed us to capture transient populations—such as LRP2.1—that do not have direct adult counterparts. Several of these developmental clusters also show spatial enrichment, being found exclusively in the insular and piriform cortices, further supporting their biological relevance.

While this pipeline was designed to accommodate diverse datasets, we observed that successful integration depends strongly on the degree of technical and experimental compatibility among datasets. In practice, datasets with large differences in sample preparation, dissociation protocol, or library chemistry often failed to meet integrability criteria, leading to exclusion from the final atlas. This limitation reflects the current sensitivity of cross-study integration rather than a restriction of our approach to in-house data.

To address this, we generated two atlas versions with differing stringency levels (Dev-SST-v1 and Dev-SST-v2; **Extended Data Fig. 6**). Dev-SST-v1 applies less stringent thresholds and includes additional external datasets, while Dev-SST-v2 retains only those meeting all three criteria, including data from the Fishell lab (Wu et al., 2024 preprint). Together, these versions illustrate both the potential and current boundaries of cross-dataset integration using our framework.

To evaluate clustering robustness and biological distinctness, we quantified cluster stability using Jaccard indices from repeated subsampling and assessed classification accuracy using random forest–based prediction of cluster identity. These analyses revealed that clusters in Dev-SST-v2 show higher stability and lower cross-talk than those in Dev-SST-v1, supporting their reliability as distinct biological subtypes.

This modular computational pipeline enables efficient extraction and analysis of any cell types from scRNA-seq datasets. While our focus here is on somatostatin-positive (SST+) inhibitory neurons, the pipeline can be seamlessly adapted to other rare populations. We demonstrated that this pipeline is applicable to brain and spinal cord datasets in both developmental and adult stages.

To maximize accessibility, the pipeline is implemented as open-source code in R and Nextflow-Conda environment, accompanied by clear documentation in Github (see Data and code availability). By combining newly generated datasets (**Extended Data Table 1**) with publicly available resources, and integrating them using our bioinformatics pipeline, we constructed the largest atlas of Sst⁺ inhibitory neurons to date. The expanded scale of the Dev-SST-v1 and Dev-SST-v2 reflect both the addition of new experimental data and the integrative computational framework that enabled consistent cross-dataset alignment. We have made this resource publicly available through a web interface (https://shiny.gbiomed.kuleuven.be/AtlasLL/), allowing users to query gene expression dynamics or subset cells by developmental stage or clusters.

### Comparison of Dev-SST-v2 to our previous work in Fisher et al. (2024)

The Dev-SST-v2 atlas extends our previous work (Fisher et al., 2024) ^23^ both in scale and analytical depth. The current dataset includes approximately sevenfold more Sst⁺ cells, providing the resolution needed to resolve transient and low-abundance subtypes. Methodologically, we replaced the MetaNeighbor-based alignment with MapMyCells, enabling more accurate, cell-by-cell cross-dataset integration and subtype identification. We also incorporated the adult reference dataset from Yao et al. (2023) ^15^— which encompasses both cortical and extra-cortical interneuron types — allowing classification beyond cortical subtypes alone. Interestingly, this expanded framework using MapMyCells initially annotated the LRP2.1 population as striatal interneurons. Upon further examination, we found that this apparent misclassification reflected the transient molecular similarity of these cortical cells to striatal interneurons, revealing that LRP2.1 represents a transient cortical population whose developmental trajectory was previously obscured. Together, these advances illustrate how increased dataset scale and refined computational integration can uncover new developmental dynamics, even through unexpected classification outcomes.

### Distinct modes of diversification within cortical SST+ interneurons

Using Dev-SST-v2, we examined the timing and modes of diversification of SST+ cortical interneurons and found significant differences in diversification strategies across SST+ subclasses. Two of these modes—invariant (predetermined) and expanding (stepwise) diversification—have been described previously in other neuronal types ^33^: the predetermined (invariant) mode has been reported for pyramidal neurons ^31,40^, chandelier neurons ^32^, and Meis2+/CGE-derived interneurons ^22^, whereas expanding trajectories were observed in retinal ganglion cells ^33^. To integrate these findings and conceptualize the temporal patterns observed in our dataset, we classified Sst⁺ interneuron trajectories into three diversification modes—invariant, expanding, and contracting (**Fig. 8**). Both invariant and expanding modes coexist within the Sst⁺ subclass, corresponding to Martinotti (MC) and non-Martinotti (nMC) lineages, respectively, while a contracting mode characterizes the LRP population, which transitions through a transient developmental state before convergence.

**Figure 8.**
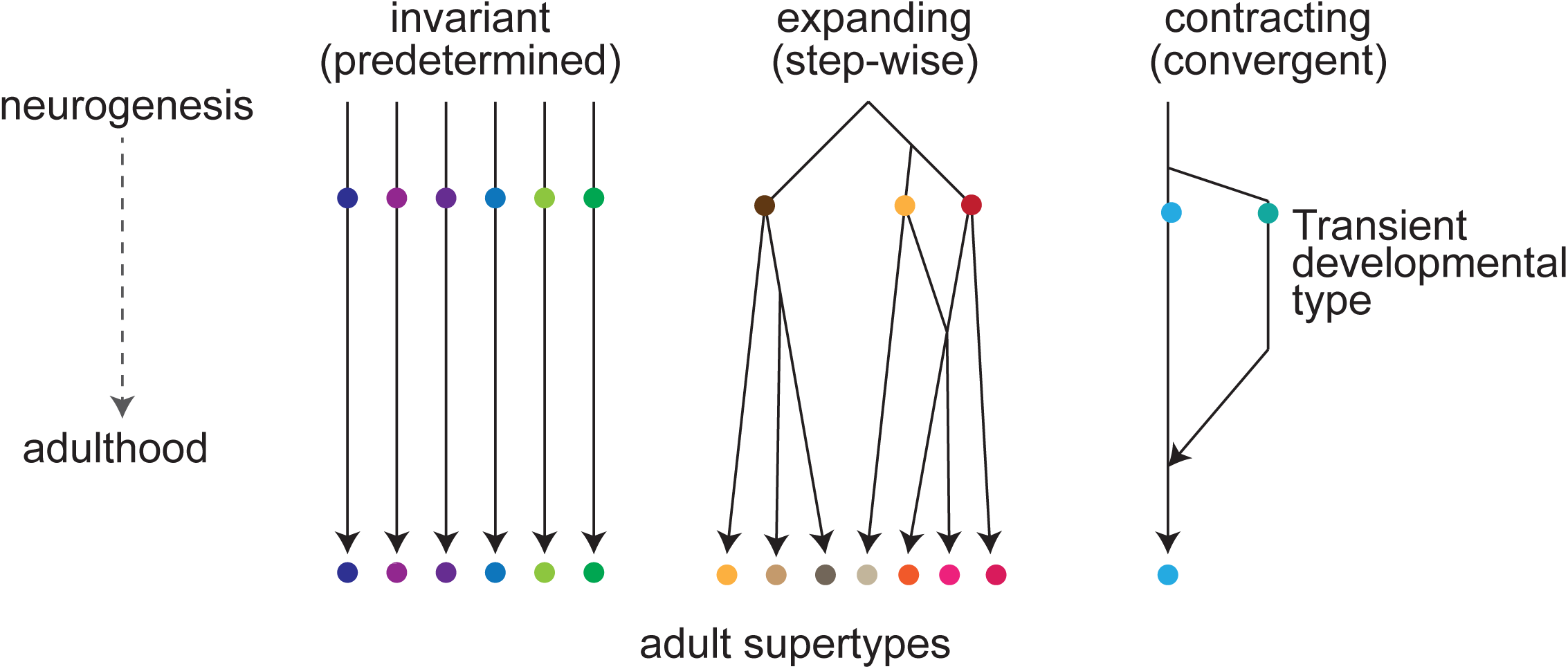
Models of diversification for postmitotic neurons. Schematic illustrating three modes of postmitotic diversification observed among cortical Sst⁺ neuron lineages. Left: Invariant (predetermined) diversification, as seen in Martinotti cells (MCs), where developmental trajectories are fixed early and cell identities remain stable from neurogenesis to adulthood. Middle: Expanding (stepwise) diversification, characteristic of non-Martinotti cells (nMCs), where successive transcriptional diversification generates new subtypes over time. Right: Contracting (convergent) diversification, exemplified by LRP subtypes, in which transient developmental clusters converge into a single mature identity. These conceptual models were adapted and modified from Shekhar et al. (2022) to illustrate the distinct temporal modes of subtype diversification revealed by the Dev-SST-v2 atlas.

For MCs, developmental diversification appears to be largely predetermined—likely established embryonically and maintained postnatally—such that early developmental clusters map directly onto adult supertypes. Many MC subtypes emerge early and remain stable, reinforcing the idea of a developmentally pre-specified trajectory. Conversely, nMCs exhibited a more gradual, stepwise diversification, with many developmental clusters mapping to multiple adult supertypes, reflecting postmitotic plasticity. Consistent with a recent preprint by the Allen Institute, Gao et al. ^54^, we found that some SST+ supertypes (e.g., 226, 227, 229) only emerge later in development. Moreover, only 3 out of 19 SST+ supertypes (0215, 0219, and 0228) stably maintain their relative abundance from P1 to adulthood, highlighting the dynamic nature of SST+ interneuron maturation.

An intriguing implication of our findings is that nMC neurons retain the capacity to alter their identity post mitotically. Supporting this idea of fluid model of cell fate acquisition, a recent preprint by Alex Pollen’s laboratory reports that a fraction of mouse SST+ interneurons, when transplanted in human organoids, begin to express Pvalb ^78^. Considering that some of the developing nMCs map to adult Pvalb+ cells, we speculate that these post-mitotic populations are intrinsically amenable in their identity.

### A convergent model of diversification in LRP neurons

One striking observation in our study is the unique ‘contracting’ mode of diversification observed in LRPs (**Fig. 8**). LRP1 and LRP2.1 are transcriptionally distinct clusters that diverge into separate developmental trajectories as early as P5. Our re-analysis of datasets ^68^ from Bax cKO mice demonstrates that LRP2.1 is selectively lost from P7 onwards due to programmed cell death.

Programmed cell death is a general developmental hallmark of cortical inhibitory neurons ^62–67^. This form of cell pruning was thought until now to target inhibitory neurons indiscriminately. We show here that programmed cell death can also target specific inhibitory populations with exquisite precision, providing a powerful mechanism for fine sculpting of cortical circuits.

Moreover, this mechanism appears to be conserved in humans. In the developing human cortex, LRP2.1 peaks at GW18–20 and is eliminated by birth, coinciding with the completion of programmed cell death in the central nervous system ^72^. The parallel trends in mice and humans suggest that selective pruning of LRP2.1 is an evolutionarily conserved strategy to fine tune the repertoire of inhibitory neurons. Collectively, our findings challenge the conventional view that inhibitory neurons follow a unidirectional and unimodal diversification trajectory by demonstrating the existence of a transient population, LRP2.1, that is selectively eliminated during postnatal cortical development.

### Implications of transient LRP2.1 neurons

The transient nature of LRP2.1 raises questions about its function during early cortical development. Transient neuronal populations, such as subplate neurons ^79^, Cajal-Retzius cells ^80,81^, and early (first wave) oligodendrocytes ^82^, are known to influence brain development by shaping circuits and connectivity. ^80,83–88^. ^6,73^. Prior studies indicate that from P4 to P6, SST+ neurons in L5b receive short-lived excitatory inputs from thalamic projections, which disappear by P7 ^6,73^. We show that similar to L5b SST+ interneurons, LRP2.1 neurons also receive transient thalamic input.

The recedes of this transient connectivity has been attributed to the metabotropic-mediated transcription, specifically the increased expression of mGluR1 in SST+ L5b neurons during the 1st postnatal ^89^. Our data suggest that another way to eliminate this transient cell type could be via programmed cell death. At the transcript level, we noted that LPR2.1, compared to LPR1, does not express high levels of mGluR1 (*Gm1*, **Extended Data Fig. 7e**), strongly implying that while these cell types have thalamic connections at P4-P6, these connections will likely not recede via an increase of expression mGluR1, but rather can be pruned off by programmed cell death. The programmed elimination of LRP2.1 by P7 and the consequent loss of these transient synaptic connections, coincides with the emergence of stable thalamic projections to L4. Future studies examining the pre-synaptic partners of LRP2.1 that could not undergo programmed cell death in adulthood could provide deeper insights into the exact function of this transient neuronal population.

In conclusion, our study presents a modular scRNAseq computational pipeline for enriching and integrating rare neuronal populations, leading to the construction of the most comprehensive developmental atlas of SST+ cortical interneurons to date. We reveal three distinct modes of neuronal diversification—invariant, expanding, and contracting. Notably, we describe a unique contracting trajectory in long-range projecting neurons, where early diversity is downscaled through selective programmed cell death. Additionally, we identified a transient cell type, termed the LIM (**<u>L</u>**ong-range, **<u>I</u>**nhibitory, **<u>M</u>**inority population) neuron, which is evolutionarily conserved. These findings provide a data-rich foundational basis for understanding how neuronal diversity is generated and refined during the development of the mammalian cortex

## Methods

### Mouse models

The mouse lines used in this study are: WT C57BL/6J (Strain #:000664, RRID:IMSR_JAX:000664), *Sst^Cre^* [B6N.Cg-Sst^tm2.1(cre)Zjh/J^] (Strain#:018973, IMSR_JAX:018973) ^50^, *RCE* [(B6.Gt(ROSA)26Sor^tm1.1(CAGEGFP)Fsh^] (Strain#: 032037, RRID:MMRRC_032037-JAX) ^90^, *RC:FLTG* [B6.Cg Gt(ROSA)26Sor^tm1.3(CAG-tdTomato,-EGFP)Pjen/J^] (Strain#:026932, RRID:IMSR_JAX:026932) ^91^ and *Nkx2.1^flp^* [Nkx2-1^tm2.1(flpo)Zjh/J^] (Strain#:028577, RRID:IMSR_JAX:028577) ^92^. All adult mice were housed in groups and kept on a reverse light/dark cycle (12/12 h) regardless of genotype. Only time-mated pregnant female mice were housed individually. Male and female mice were used in all experiments. SST+ cells labelled by *Sst^Cre^* mice were crossed with RCE reporter mice containing a floxed-stop cassette upstream of enhanced green fluorescent protein (*EGFP*) in the ROSA gene locus to obtain *Sst^Cre^;RCE^fl/+^*. In some experiments, we isolated MGE-derived cells, by using *Nkx2.1^flp^; RC:FLTG*. All mice were maintained on a C57Bl/6 background and housed under a 12 h light/dark cycle with ad libitum access to food and water. All procedures were approved by KU Leuven and Scripps animal welfare committees.

### Tissue preparation for FACS

Tissue preparation was similar to Fisher et al., (2024) ^23^. These include samples from DS19, DS20, DS21, DS22, DS23, DS24 and DS26. Briefly, all solutions used in this procedure have an osmolarity of 290 ± 20 mOsm/kg. For E16.5 and P1, tissue from 3 animals was pooled together. For P5, tissue from 1 and 2 animals was pooled together. The neocortex of E16.5, P1 and P5 *SST^cre/+^;RCE* mice were dissected in an ice-cold HBSS buffer supplement with 25 mM of glucose. Tissue was cut into small pieces and digested for 10 min at 34°C with carbogen oxygenation using 30ml of dissociation buffer solution, pH7.5, with the following composition: 0.2 mg/ml pronaseE (Sigma), 50 mM of trehalose, 10 mM glucose, 0.8 mM kynurenic acid, 0.05 mM APV, 0.09 M Na_2_SO_4_, 0.03 M K_2_SO_4_ and 0.014 M MgCl_2_. After 10 mins of incubation, E16.5 tissue was incubated at 34° for 5 more minutes with 375 µl of 10mg/ml DNAse while P1 and P5 tissue was incubated for 5 more minutes with 375 µl of 10mg/ml DNAse and 750µl of 10mg/ml liberase (Sigma). Post enzymatic digestion, tissues were washed once in an ice-cold dissociation buffer without enzyme. Tissue was mechanically triturated into single cells in 0.5 ml (for E16.5) or 5ml (for P1 and P5) of OptiMEM (Invitrogen) supplement with 2% Trehalose, 10 mM glucose, 0.8 mM kynurenic acid and 0.05 mM APV using a volumetric pipette. For P1 and P5 tissue, myelin was removed by overlaying single cell suspension on top of a 20% percoll and centrifuge at 700 g for 10 mins at 4 °C. The cell pellet was resuspended in 400 ul of OptiMEM with 2% Trehalose, 10 mM glucose, 0.8 mM kynurenic acid and 0.05 mM APV and tissue chunks were eliminated by passing cells suspension through a 40 mm cell strainer (BD).

For sample DS18 only, pregnant C56BL/6 dams carrying E18.5 embryos, previously injected intraventricularly with AAV expressing eYFP, were anesthetized with isoflurane and sacrificed by decapitation. The uterus was removed, the embryos were moved into cold Leibovitz’s L15 medium supplemented with 7 mM HEPES, and each embryo was decapitated. From each brain a section of the neocortex, corresponding roughly to S1, was dissected out with a micro-knife, avoiding the underlying striatal tissue. Tissue from 4-5 embryos of the same litter was pooled. Tissue sections were dissociated into single cell suspensions using the Worthington Tissue Dissociation Kit; the papain, DNAse, and albumin ovomucoid inhibitor solutions were reconstituted and oxygenated according to the manufacturer’s instructions. Tissue was digested in papain/DNAse for 15 min at 37° with carbogen oxygenation, triturated in EBSS using 1 mL lo-bind pipette tips, and then the cell suspension was mixed with albumin ovomucoid inhibitor/DNAse and centrifuged at 230g for 5min in a swinging bucket centrifuge. Cells were resuspended in an OptiMEM-trehalose media (OptiMEM supplemented with 50 mM of trehalose, 10 mMglucose, 0.5 mM kynurenic acid, 0.025 mM APV, 4 mM MgCl_2_, 0.2 mM HEPES). Debris was removed by overlaying the suspension on top of a 5% Optiprep gradient and centrifuging at 100g for 10 mins. The cell pellet was resuspended in 500 uL of optimem-trehalose media, and DAPI and DRAQ5 were added to a final concentration of 1uM.

For dataset DS23, after myelin removal the cell pellet was resuspended in 1ml of 3% Glyoxal with the following composition: 2.8 ml Nuclease Free H_2_0, 0.79 ml 100%EtOH, 0.31ml 40% Glyoxal (Sigma 50649), 30ul acetic acid, ph 4-5, for 15 min at 4° shaking. After incubation, cells were centrifuged at 800g for 10m at 4° and resuspended in 500ul of 1X PBS supplemented with 1% BSA and tissue chunks were eliminated by passing cells suspension through a 40 mm cell strainer (BD).

### Cell isolation and 10X Genomics scRNAseq

GFP+ and DAPI negative for live samples and GFP+ and Fixable Viability Dye eFluor 780 negative cells for fixed samples were isolated using fluorescence-activated cell sorting (FACS) with a flow cytometer (BD Aria Fusion) using a 100 µm nozzle, about 3000-5000 events per second, at purity mode, and collected directly in 600 µl (cold 4 deg) of OptiMEM with 2% Trehalose, 10 mM glucose, 0.8 mM kynurenic acid and 0.05 mM APV solution in a 1.5-ml LoBind (Eppendorf) tubes precoated with 3% BSA overnight. We limited the FACS procedure to a maximum sort time of 25 mins to ensure high cell survival. We collected 20,000 to 30,000 cells for each sample. We centrifuge the collect cells at 600g for 5 minutes at 4° and resuspend the FACS cells in 40 µl of OptiMEM with 2% Trehalose, 10 mM glucose, 0.8 mM kynurenic acid and 0.05 mM APV. To determine cell number and viability, 5 µl of Trypan Blue was added to 5 µl of cells and cells were counted on a hemocytometer. In all experiments, we only proceed toward scRNA-seq with > 80% viability. About 12000 isolated single cells for each sample were loaded onto a custom droplet encapsulation platform, HyDrop ^93^, for single-cell capture. cDNA library preparation was performed following the 10X Genomic commercial kit CG000315. RNA-seq was performed using BGI MGISEQ-2000 and Illumina NovaSeq 6000 v1.5.

For sample DS18, cell suspension was sorted for live eYFP+ cells using an Astrios EQ sorting flow cytometer with a 100 um nozzle at 2-way purity. Sorted cells were collected in 0.5 ml of cold FACS media (OptiMEM supplemented with 50mM of trehalose, 10mMglucose, 0.5 mM kynurenic acid, 0.025 mM APV, 0.2 mM HEPES). Approximately 20,000 cells were collected. Cells were centrifuged at 200g for 15 minutes at 4° in a swinging bucket centrifuge and resuspended them in about 20 µL of FACS media. 5 µL of the cell suspension was taken out for cell counting and viability with a Countess, and we only used suspensions with >80% live cells. About 9000 isolated single cells for each sample were loaded onto a 10X Genomic single-cell chip for GEM formation and cDNA library preparation. RNA-seq was performed using an Illumina Nextseq200 P2 flow cell.

### Shallow and deep scRNAseq

cDNA libraries were sequenced at two different depths. For shallow sequencing, we targeted a sequencing depth of 5,000 reads per cell; the actual achieved read number was 8,227.5 +/- 793.5 (mean +/-s.d.). For deep sequencing, we targeted 50,000 reads per cell; the actual read number achieved was 61,384 +/-20,484.

### Data pre-processing

Sequencing data were prepared for analysis by application of the 10X CellRanger pipeline. First, Illumina and BGI BCL output files were de-multiplexed into FASTQ format files. Features counts were computed for individual GEM wells. STAR aligner ^94^ was applied to perform splicing aware alignment to the mm10 reference genome, and then reads were bucketed into exonic, intronic, and intergenic categories. Reads were classified as confidently aligned if they corresponded to a single gene annotation. Only confidently aligned reads were carried forward to UMI counting. A cell calling algorithm in conjunction with the barcode rank plot was applied to filter low RNA content cells and remove empty droplets. The output is a read count matrix. For publicly available datasets we downloaded raw count matrices. We used the R package Seurat (v 4.3.0.1) to perform the bulk of our subsequent data analysis, including filtering, normalization, scaling, and other downstream processing.

### Sample processing and filtering of Sst+ cells (Module 1)

Each sample has been first processed separately for cell filtering and identification of cortical Sst+ neurons. We discarded any genes detected in fewer than 10 cells before filtering low quality cells. Cells were retained if they met the following criteria: (i) unique gene count > 700, (ii) mitochondrial gene content < 10%, (iii) ln(total UMI) > mean(ln(total UMI))-2sd(ln(total UMI)). To assess cell cycle phase differences in our dataset, we utilized the *CellCycleScoring* function from Seurat. Cell cycle phase scores were calculated for each cell using predefined gene lists for the S phase and G2M phase. Normalization and scaling of reads were performed using the R package Seurat, with the functions *NormalizeData()* and *ScaleData()*. During scaling, the number of total genes, percentage of mitochondrial gene expression and cell cycle phase difference score were regressed out to remove unwanted sources of variation from the data. Highly variable genes were defined using the Seurat function *FindVariableFeatures()* and Principal Component Analysis (PCA) was performed with the function *RunPCA()*. Doublets were identified using the scDblFinder ^95^ (v 1.12.0) algorithm with doublet rate set to 0.07. Cell clustering annotations were provided to scDblFinder, derived by applying *TransferData()* function to the cluster labels of our previously published dataset ^23^ using anchors identified by *FindTransferAnchors()* function on top 40 principal components. The optimal number of principal components was estimated performing a cross-validation on Principal Component Analysis (PCA) ^96^ and the resulting number of components was used as input to compute a k-Nearest Neighbors (k-NN) graph further refined into a Shared Nearest Neighbor (SNN) by the Seurat function *FindNeighbors()*. Clusters were defined using the Louvain algorithm implemented in *FindClusters()* function. Uniform manifold approximation and projection (UMAP) embedding was also computed running *RunUMAP()* with the estimated optimal principal components. We selected cells belonging to clusters expressing Sst gene, discarding “stressed” clusters, characterized by high expression of heat shock genes and lower average UMI, and clusters expressing Meis2, QC metric similar to Fisher et al., 2024 ^23^.

### Integrability testing of datasets (Module 2)

To test if any given dataset can be used to generate an integrative atlas, we selected only datasets containing at least 1,000 high-quality Sst+ cells, as identified using previously described methods. The integrability of each dataset with our reference atlas (Atlas-v0) was evaluated using three criteria: Anchor points, Neighborhood composition, and kBET ^52^ rejection test.

1. **Anchor Points** -The integration anchors between the testing datasets and Atlas-v0 were identified using the *FindIntegrationAnchors()* function in the Seurat package, with reduction = ‘cca’. For each dataset, we subset 1000 cells and compute the mean number of anchors for each testing dataset relative to Dev-SST-v0. This is termed as a normalized mean anchor point as we always subset 1000 cells. A normalized mean anchor count greater than 1686 was set as the acceptance criterion.

2. **Neighborhood composition** — The testing dataset was integrated with Atlas-v0 using the **IntegrateData()** function based on previously identified anchors to generate a corrected gene expression matrix. This matrix was scaled using **ScaleData()**, and principal components were computed using **RunPCA()**. Pairwise Euclidean distances were then calculated between cells in the testing dataset and cells in the reference atlas (Dev-SST-v0) within the integrated PCA space. For each test cell, the minimum distance to reference cells was recorded and added as metadata. k-NN Computation: Nearest neighbors were identified using the *dbscan::kNN()* function in PCA space. For each k-value, the proportion of cells meeting the threshold of reference atlas cell representation was computed.

To evaluate neighborhood composition, we calculated the proportion of test dataset cells whose nearest neighbors contained at least half the expected fraction of reference atlas cells across a range of *k*-values (5–200, in increments of 5). Datasets in which at least 50% of test cells had at least half the global fraction of reference atlas cells among their *k* = 60 nearest neighbors were considered to meet the integration criteria. To assess the similarity of local transcriptomic structure between datasets, we constructed a *k*-nearest neighbor (kNN) graph for each dataset (*k* = 60). Two datasets were considered to have similar neighborhoods when at least 50% of the *k* = 60 nearest neighbors of corresponding cells overlapped.

3. **kBET Rejection Rate** - The kBET ^52^ rejection rate (version 0.99.6) was calculated to evaluate how well the testing dataset and reference atlas (Atlas-v0) were integrated in the corrected PCA space. A rejection rate below 50% was set as the acceptance criterion.

Only datasets meeting all three criteria (anchor points, neighborhood composition, and kBET rejection rate) were included in the generation of Atlas-v1.

### Data Integration and annotation (Module 3)

Individual samples were integrated into a single object using canonical correlation analysis (CCA) ^97^. Anchors were identified across samples with *FindIntegrationAnchors()* function with parameters reduction= ‘cca’, k.filter=100, k.score=15 and anchor.features=2000. A batch corrected gene expression matrix was then computed using *IntegrateData()* function. The resulting matrix was scaled using *ScaleData()* function, regressing for number of total genes, percentage of mitochondrial gene expression and cell cycle score. PCA was performed with *RunPCA()* function, and the optimal number of principal components was determined via cross-validation. Clustering was computed with *FindNeighbors()* and *FindClusters()* functions, based on the optimal number of principal components estimated. Uniform manifold approximation and projection (UMAP) embedding was computed using *RunUMAP()* function. Cells within “stressed” clusters characterized by high expression of heat shock genes and low average UMI, doublets, or clusters showing high expression of mitochondrial genes were further removed. Resulting cells were annotated to major groups of cortical Sst+ neurons (LRP, MC, and nMC) using annotated cells from the base atlas. This annotation was achieved by applying the *TransferData()* function on anchors identified between each sample and the base atlas using the *FindTransferAnchors()* function. Each cell was annotated to the group that achieved the maximum score.

### Iterative Clustering of Dev-SST-v0, v1, v2

Clustering algorithm was performed separately on each major group of cortical Sst+ neurons (LRP, MC and nMC). To compute clusters, we followed an iterative approach characterized by the following steps:

1. Sample filtering. Remove samples with less than 20 cells.
2. Data integration and clustering. Perform data integration using canonical correlation analysis and apply clustering as previously described. Clustering was computed at resolutions of 0.1, 0.2, 0.3, 0.4 and 0.5 using a kNN graph built with k=20 using the optimal number of principal components (PCs). Duplicated clustering (clusters from different resolutions that result in the same cell annotation) were considered only once.
3. Clustering stability evaluation. We performed 20 data sub-samplings, each time sampling 80% of the cells without replacement, and repeated the clustering procedure for all selected resolutions on each subsample. Jaccard indices were calculated for each cluster at each resolution by comparing the original clusters with those obtained from each subsample. We defined a stable cluster as one with Jaccard index 96 greater than 0.75 in at least 75% of the subsamples. A stable set of clusters was defined as one where more than 70% of the cells belonged to stable clusters and stable clusters accounted for more than 60% of the total clusters. These parameters were selected based on recommendations from Tang et. al ^98^. If no clustering was found to be stable, the iteration was stopped, and the cluster label obtained in the previous iteration was assigned as the final identity of all the cells. Otherwise, among stable clustering, we selected the resolution that produced the higher percentage of cells within stable clusters. If two or more clustering resulted in the same percentage, the resolution with the higher number of clusters was chosen.
4. Cluster identity evaluation and merging. Clusters with fewer than 300 for Martinotti and Non-Martinotti and 150 cells for LRPs were merged with their closest cluster: centroids were calculated in batch corrected PCA space and the closest cluster was defined as the one with the smallest Euclidean distance between centroids. For each resulting cluster, marker genes were identified by comparing gene expression with all other cells not belonging to the cluster. FindMarkers() function from Seurat was used with parameters logfc.threshold=1 and only.pos = TRUE. Genes with an adjusted p.value lower than 0.01 were selected and the sum of the min (-log10(p.val_adj), 20) values across all selected genes was calculated (DEScore). If the resulting value was less or equal to 60, the cluster was merged with its closest cluster as previously described. The cutoff of DEScore >60 was retained from our previous work (Fisher et al. 2024) ^23^.
5. Process iteration. After identity evaluation, if only one cluster remained, the iteration was stopped and the cluster label obtained in the previous iteration was assigned as the final identity of the cluster. If more than one cluster remained, the process was repeated from step 1 to step 5 for all clusters with 600 cells or more for Martinotti and Non-Martinotti; and 300 for LRPs. Clusters with fewer than 600/300 cells were returned as the final identity.
6. After all clustering iterations ended, cells removed at step 1 of each iteration were annotated. The k-NN constructed during iteration 1 was used to identify the nearest neighbours for each removed cell, and each cell was assigned to the cluster most represented among its nearest neighbours. Finally, we computed marker genes for all clusters using FindAllMarkers() function from Seurat with parameters logfc.threshold=log2(1.5), min.pct=0.2 and only.pos = TRUE. Based on the resulting gene list we build a hierarchical clustering dendrogram of clusters using Euclidean distance. We then identified the 5th percentile of the distribution of cophenetic distances across all cluster pairs, where the cophenetic distance is defined as the height of the dendrogram at which the branches containing two clusters merge. For all pairs of clusters with cophenetic distance below this threshold, we compute the DEScore, as previously defined, in a one vs one cluster comparison. If the two clusters had a DEScore less or equal to 60, they were merged into a single cluster.

### Random Forest Classification and Atlas Comparison

To evaluate and compare the performance of two independently generated single-cell RNA-seq atlases (Dev-SST-v1 and Dev-SST-v2), we trained random forest classifiers to predict cell-type identity based on gene expression profiles. Classification models were constructed separately for each major interneuron family (LRPs, Martinotti, and Non-Martinotti). For every cell type within each family, we computed confusion matrices quantifying how frequently each true class was assigned to each predicted class.

From these matrices, we extracted per-class performance metrics: Diagonal accuracy is defined as the fraction of cells correctly predicted for a given class (true-positive rate per class). Cross-talk is defined as [1 - (diagonal accuracy)], representing the degree of misclassification to other classes. For each atlas version (v1 and v2), we summarized these metrics and computed per-class differences (Δ= v2 - v1) for both diagonal accuracy and cross-talk. Positive Δ (diagonal) values indicate improved classification accuracy in v2, while negative Δ (cross-talk) values indicate reduced confusion between classes.

### Cell annotation validation

To validate cell identities, we performed five-fold cross-validation, taking 20% of the data as a test set and the remaining 80% as the training set. We computed differentially expressed genes for all clusters in the five training sets using *FindMarkers()* function with options logfc.threshold=ln(1.5), only.pos=TRUE, min.pct=0.5. We then proceed in a pairwise fashion, using *randomForest* ^99^ R package (v 4.7-1.1) we trained a random forest classifier with 100 trees on each pair of clusters in the training data using only the top 10 differentially expressed genes, ranked by logFC, from each cluster. This classifier was used to predict the identity of cells in the test set. Following five-fold cross-validation, we have a predicted identity for every cell for each pairwise comparison of clusters. The above procedure was repeated for a total of 10 iterations. From the consistency of the outcome across these iterations, we extracted a measure of the stability of the cell classification. We used cell_type_mapper, the backend python package of the MapMyCell tool, to perform cell type annotation on custom datasets.

### Waddington-Optimal Transport analysis

We performed Waddington-Optimal Transport (WOT) ^100^ analysis to infer lineage relationships and cell fate transitions across developmental time points in the major subgroups of cortical Sst+ neurons (LRP, MC, and nMC). All analyses were conducted using the wot command-line interface (v. 1.0.8). For both Martinotti and non-Martinotti subsets, cluster sizes were imbalanced. To address this, we computed the median cluster size across all clusters and randomly subsampled this number of cells from each cluster. This approach allowed us to generate a more balanced dataset for fate prediction. The subsampling procedure was repeated three times, and WOT analysis was performed independently on each random subset. First, transport maps were generated between consecutive developmental stages using the optimal_transport function. As input, we used a log-normalized count matrix restricted to highly variable genes, to reduce noise and focus on the most informative features. The --growth_iters parameter was set to 3 to allow estimation of population growth rates across iterations, thereby improving transport map stability. Next, fate probabilities were computed with the fates function, applied to the resulting transport maps. This procedure estimated, for each cell, the probability of transitioning into a given cluster identity at subsequent time points. To facilitate interpretation, cells at each time point were grouped according to their cluster assignments, and for each cluster we calculated the median fate probabilities toward all possible cluster identities at the following time point. For Martinotti and non-Martinotti subsets mean values across the three subsampled datasets were reported.

### Cell annotation to adult counterparts

To annotate our data to their adult counterparts, we used the *MapMyCells* ^15^ algorithm, using the 10x Whole Mouse Brain (CCN20230722) as reference taxonomy and using the hierarchical mapping as chosen algorithm. All annotations of cell identity in the adult reference were used; we did not restrict the dataset by brain region or cell class. Briefly, to annotate unlabelled cells to their corresponding class, subclass, supertype, or cluster, the algorithm compares the gene expression of each unlabelled cell with the average expression profile of each child node at every level of the hierarchy. This comparison is performed using marker genes that effectively discriminate among child nodes, assigning a vote to the child node with the highest Pearson correlation. For robustness, the process is repeated 100 times, randomly selecting 90% of all marker genes every time. A cell is assigned to the child node with the highest number of votes, and cell labels were considered valid if 90 out of 100 annotations assigned the cell to the same child node. We then defined a 1:1 matching between one of our clusters and an annotated supertype if at least 65% of the cells within the cluster were assigned to that supertype, and if the majority of all cells annotated to that supertype originated from that cluster, representing at least 30% of all cells annotated to the supertype.

### Cortical and Striatal Integration

Cells sorted from striatum were processed as previously described (Module 1), resulting in the identification of high-quality Sst⁺ cells. These cells were mapped to the Whole Mouse Brain reference (CCN20230722) using the MapMyCell tool, as described earlier. In addition, Seurat label transfer analysis was performed to map cells to cortical atlas v1 clusters, using the FindTransferAnchors() function followed by the TransferData() function. Over 90% of cells were assigned to the LRP2 cluster. We selected these cells and integrated them with cortical LRP2 cells from the same developmental stage (P5). To achieve soft batch correction while preserving biological differences between samples of different origins, we performed FastMNN analysis ^101^. Specifically, the RunFastMNN() function from SeuratWrappers package (v. 0.3.1) was applied to Seurat objects containing LRP2 cells, split by sample_id. A UMAP was then computed on the top MNN reduction to visualize the cells. Differentially expressed genes between the two datasets were identified using the FindMarkers() function, with parameters min.pct = 0.2 and logfc.threshold = 1.

### Trajectory and Pseudotime analysis

We performed trajectory analysis on the LRP clusters. First, we computed gene modules that captured distinct sources of variation within the data. We used the R package Antler ^56^ (v 0.9.0) to generate these modules. Computation began with a normalized counts-per-million matrix. Genes present in fewer than 10 cells were discounted from the computation. Genes were filtered prior to clustering to retain only those genes having Spearman correlation with at least 5 other genes at value 0.3. These genes were clustered hierarchically using Spearman correlation as a dissimilarity metric. This hierarchical clustering was repeated over many iterations, with a heuristic algorithm determining an optimal number of modules. At each iteration, modules were discarded if they failed to meet validity thresholds on the expression level of individual genes. At the terminal iteration, all modules met the validation criteria and were returned as the final set of gene modules. To understand the module contents, we performed functional enrichment on each module and examined gene ontology terms. We selected 335 genes from modules associated with terms including “neuron differentiation”, “axon development”, and “synapse assembly”. We computed Diffusion Maps with Destiny ^102^ R package (v 3.12.0) on the corrected expression matrix, considering only the selected genes. The eigenvalues of the diffusion map transition matrix were used as diffusion components for low-dimensional embedding. The pseudotime for each cell was computed from the full transition matrix using the *DPT()* function and represents a notion of diffusion distance from a root cell. The root cell was identified as the cell from the earliest age samples that had the highest number of early sample cells in its neighborhood (computed on diffusion components). From this analysis, we inferred linear branches diverging from an initial shared state. We then identified genes exhibiting continuous changes over pseudotime, by fitting each gene’s expression vector to pseudotime using a generalized additive model (GAM), with the *gam()* function from *mgcv* ^103^ R package (1.9-0). Genes were classified as early genes (negatively correlated with pseudotime) or late genes (positively correlated with pseudotime). Late genes were further subdivided into those specifically expressed in only one branch.

### Saturation analysis

We subsampled cells without replacement at percentages ranging from 5% to 95% in 5% intervals. For each percentage, we generated 10 different subsamples. To estimate the saturation point, we calculated several metrics. First, we computed robust Hausdorff Distance between each subsample and the original dataset in the batch corrected PCA space, selecting the 50th largest value as the distance measure. Next, we identified marker genes for each cluster in each subsample and recorded the total number of marker genes. Additionally, we calculated the absolute differences in mean expression for marker genes between the subsamples and the entire atlas for each cluster. The mean of differences across clusters were summed over all differentially expressed genes. Finally, we assessed the fraction of clusters that maintained their identity in each subsample. For this, cluster centroids were calculated based on gene expression vectors, and clusters from the whole dataset were matched to the centroids of the subsamples using *MetaNeighborUS()* function from MetaNeighbor ^55^ R package (v 1.18.0). Clusters with an AUROC score greater than 0.9 when matched to their centroids in the subsample were considered to have retained their identity. For each subsampling percentage, we computed the mean scores for all metrics (robust Hausdorff Distance, number of differentially expressed genes, sum of absolute differences in mean expression, and percentage of clusters preserving their identity) across the 10 subsamples. The inflection point, indicating the saturation percentage, was identified as the point where the second derivative of the metrics changed sign.

### Cloning of viral vector and generation AAV

The AAV enhancer plasmid, pAAV-eHGT-m600-sYFP2 ^60^, was provided by Boaz Levi (Allen Institute). To generate pAAV-eHGT-m600-dio-sYFP2, sYFP2 was removed with restriction enzymes BamH1 and EcoRV, and the backbone was isolated by gel purification. The insert, dio-sYFP2, with a 20-bp homology on both 5’- and 3’-end to the backbone was synthesized by Twist Bioscience as a gene fragment. Backbone and insert were assembled by NEBuilder Hi-Fi DNA Assembly (NEB, E2621L) per manufacturer recommendation and transformed into NEB Stable competent cells (NEB, C3040I). Using the transfer vectors pAAV-eHGT-600m-sYFP2 and pAAV-eHGT-600m-dio-sYFP2, AAV2/9 were custom generated by BrainVTA at a viral titre of >3 x 10^13^ vg/ml.

### Neonatal injection of AAV

AAV injections were performed on P0 wild-type CD1 mice or *SST^Cre^*^/+^ mice under hypothermia-induced anesthesia. Injections into the lateral ventricle were conducted using glass micropipettes (1B120F-4, World Precision Instruments) connected to a pressure microinjector device (µPUMP, World Precision Instruments). The injection site was identified at half the distance from the lambda suture to each eye. The glass micropipette was inserted perpendicularly into the skull to a depth of approximately 2 mm, with slight resistance decrease indicating entry into the lateral ventricle. Approximately 1 µL of the virus (AAV2/9-eHGT-600m-SYFP2 or AAV2/9-eHGT-600m-dio-SYFP2) was rapidly injected into each ventricle. Following the injections, pups were placed on a warming pad to recover until they could move normally and were then returned to their mother.

### Neonatal injection of AAV and Rabies virus

Rabies helper AAVs containing a cre dependent GFP, TVA, and optimized G protein (Flex-nGToG) were injected into the lateral ventricles of *Sst^cre^* mice at postnatal day 0 (P0). Each brain received 2 µL of helper virus (final titer 5 × 10¹² vg/ml). Two helper viruses were used to generate starter cells from either all SST⁺ neurons or long-range projecting (LRP) SST⁺ neurons. For pan-SST⁺ targeting, pAAV-syn-Flex-nGToG (titer > 1 × 10¹³ vg/ml) was obtained from the Charité Vector Core (Berlin, Germany). For LRP-restricted targeting, pAAV-600m-Flex-nGToG was subcloned by PCR from pAAV-syn-Flex-nGToG using pAAV-eHGT-600m-dio-sYFP2 as the backbone. AAV2/9 particles were custom produced by VectorBuilder (USA) at titers > 1 × 10¹³ vg/ml. At P4, the rabies pseudovirus N2c-ΔG-Tdt-Flp (Charité Vector Core; titer = 1.06 × 10^9^ IU/ ml) was injected into the somatosensory cortex using a pulled glass capillary and a nanoinjector (NanoLiter 2020, WPI). Three injections of 50 nl each were delivered at 25 nl per min. Injection sites were defined relative to the intersection of the superior sagittal and transverse sinuses: Injection 1: DL = 3.1 mm, ML = 2.2 mm, depth = 0.6 mm; Injection 2: DL = 3.3 mm, ML = 2.1 mm, depth = 0.6 mm; Injection 3: DL = 3.5 mm, ML = 2.0 mm, depth = 0.6 mm. Injection accuracy was verified by co-injection of fluorescent beads and post hoc histological validation. Starter cells (GFP+ and Tdtomato+ cells) and presynaptic connected cells (Tdtomato+ cells) are quantified by image segmentation script written in Python.

### Rabies quantification and statistics

Three brains per starter line (LRP and panSST+) were analyzed, comprising a total of 54 sections (9 sections per brain). For each brain, the number of Red⁺ cells per mm² in each ROI was normalized to 100 starter cells and averaged across all sections to yield section-level mean densities per region and starter type. Differences between starter lines were assessed using linear mixed-effects models with starter type as a fixed factor and brain identity as a random intercept to account for repeated sections. Separate models were fit for each anatomical region and for each outcome (normalized Red⁺ density and Double⁺ density). P values were adjusted for multiple comparisons using the Holm–Bonferroni method (α = 0.05). Section-level data are shown as gray points overlaid on boxplots (median ± IQR) with light pink (LRP) and blue (allSST) fills. Analyses were performed in Python (statsmodels, matplotlib).

### Histology and immunochemistry

Adult and postnatal pups were deeply anaesthetized with an intraperitoneal injection of ketamine (87 mg per kg body weight) and xylazine (13 mg per kg body weight) and transcardially perfused with ice-cold phosphate-buffered saline (PBS) followed by 4% paraformaldehyde (PFA). The brains were isolated and incubated overnight in 4% PFA on a slow-speed rocking platform at 4 °C. Only for VgluT2 staining, postnatal pups were transcardially perfused with ice-cold phosphate-buffered saline (PBS) followed by 9% glyoxal, 8% acetic acid solution (pH4) as a fixative. Tissue was washed with PBS, incubated in 10% followed by 30% sucrose in PBS and cut on a freezing microtome into 100 μm coronal sections, and stored in cryoprotectant solution (30% glycerol, 30% ethylene glycol in PBS) at -20 °C or processed or free-floating immunohistochemistry. Coronal sections were stained using the following primary antibodies: rabbit anti-Dach1 (1:500, Proteintech, 10914-1-AP), chicken anti-GFP (1:3000, Aves Lab, GFP-1020), rabbit anti-MafB (1:500, Sigma Aldrich, HPA005653), rabbit anti-Nitric Oxide Synthase 1 - NOS1 (1:1000, Immunostar, 24287), and guinea pig anti-VgluT2 (1:1000, Merck, AB2251). Tissue was blocked with 2% BSA, 10% goat serum in 0.25% triton-X-100-PBS for 1h at RT. Primary antibodies were incubated overnight at 4 °C in 2% BSA, 10% goat serum in 0.25% triton-X-100-PBS. The following secondary antibodies conjugated to Alex Fluor dyes were used: goat anti-chicken IgY (H+L) Alexaplus 488 (Thermofisher A32931), and donkey anti-rabbit IgG (H+L) Alexaplus 647 (Thermofisher A32795). Secondary antibodies were all used at 1:1000, incubated for 2 h at RT, followed by 4’,6-diamidino-2-fenylindool (DAPI) (5 μM, Sigma, D9542) for 10 minutes at RT. Sections were washed three times for 10 minutes with 0.1% triton-X in PBS after secondary and DAPI staining. Brain sections were mounted in Mowiol-DABCO (25% Mowiol, Sigma, 81381, 2,5% DABCO, Sigma, D27802). Brain sections were imaged using an inverted Ti2-E Nikon microscope with a Crest Optic X-light V3 spinning disk. Subsequent image processing was performed using FIJI or Matlab.

### Image Analysis and Quantification of LRP1 and LRP2.1 by histology

Two types of staining were used for the analysis: (i) GFP and Dach1, and (ii) GFP and MafB. LRP1 was defined as GFP+ cells expressing high levels of MafB but low levels of Dach1. In contrast, LRP2.1 was defined as GFP+ cells expressing low levels of MafB but high levels of Dach1. Quantification of Dach1 and MafB expression levels was performed using MATLAB with a customized script, which is publicly available on GitHub. Briefly, GFP+ cells were selected based on a manual thresholding function determined by the user. For each GFP+ cell, the centroid was identified and expanded into a circular region with a 4-pixel radius. The mean intensity signal of either MafB or Dach1 within this circular region was then calculated. For Dach1, mean intensity values in the GFP+ cells that were at least 2× above the background intensity were classified as high Dach1. For MafB, since all GFP+ cells expressed MafB, the intensity values in GFP+ cells were fitted to a normal distribution. Cells with MafB intensity levels above the 75th percentile (mean + 0.524 standard deviations) were classified as high MafB. Quantification of VgluT2 puncta area was performed using MATLAB with a customized script by thresholding the GFP channel to compute cell area; thresholding of theVglut2 staining was used to quantify VgluT2 spot area.

### Slice preparation for electrophysiology

P4–P6 mice were anesthetized by hypothermia and decapitated. Brains were rapidly dissected in ice-cold, oxygenated (95% O₂ and 5% CO₂) sucrose-based cutting solution containing (in mM): 83 NaCl, 2.5 KCl, 1 NaH₂PO₄, 26.2 NaHCO₃, 3.3 MgSO₄, 0.5 CaCl₂, 22 D-glucose, and 72 sucrose (pH 7.4). To preserve the thalamocortical pathway, brains were cut at a 55° angle relative to midline in combination with an additional 10° ramp (Agmon and Connors, 1991). Somatosensory thalamocortical slices (350 μm) were prepared (Leica VT1200) and allowed to recover in cutting solution at 32 °C for 30 min, followed by at least 30 min at room temperature before recordings. Initially tetramethylrhodamine crystals (3000 MW, Thermo Fisher Scientific) were introduced into the thalamus to guide selection of slices with intact thalamocortical pathway (e.g. Stokke et al., JCN 2002). Ultimately, optimal slices could be identified visually, eliminating the need for additional fluorescent labelling.

### Electrophysiology patch clamp recordings and analysis

Slices were transferred to a submerged chamber (Warner Instruments) and continuously perfused (3–4 ml/min) with oxygenated artificial cerebrospinal fluid (ACSF, 32-33 °C) containing (in mM): 127 NaCl, 2.5 KCl, 2.5 NaH₂PO₄, 2 CaCl₂, 2 MgCl₂, 25 NaHCO₃,25 D-glucose. Bicuculline (20 μM) was added to the ACSF in all recordings. Recording electrodes (3–6 MΩ) were pulled from borosilicate glass capillaries using a horizontal P-1000 Flaming/Brown micropipette puller (Sutter Instrument) and filled with an internal solution containing (in mM): 126 CsMeSO₃, 10 HEPES, 2.5 MgCl₂, 4 ATP, 0.4 GTP, 10 creatine phosphate, 0.6 EGTA, 5 QX-314-Cl, and 3 mg/ml biocytin, adjusted to pH 7.25 with CsOH. Data were acquired using a Multiclamp 700B amplifier (Molecular Devices), digitized at 20 kHz, and low-pass filtered at 3 kHz via a Digidata 1440A interface (Molecular Devices). pClamp 10 software (Molecular Devices) was used for acquisition. Input resistance, series resistance, and holding current were continuously monitored to ensure recording stability and data quality. Series resistance (<20 MΩ) was compensated by 70-75%. Spontaneous excitatory postsynaptic currents (sEPSCs) were recorded in whole-cell voltage-clamp mode at a holding potential of –70 mV for 3 min. For evoked EPSC recordings an extracellular stimulation electrode (cluster microelectrodes, FHC) was positioned onto the internal capsule (IC) axons exiting the ventrobasal (VB) thalamic nucleus. To ascertain activation of single axons, we first determined the minimal stimulation intensity required to successfully evoke EPSC in 50-60% of the trials (10-45V, A-M Systems Model 2100).This voltage was subsequently used for evoked recordings. Evoked EPSCs were analyzed using Easy Electrophysiology, and spontaneous EPSC was analysed using Mini Analysis software (Synaptosoft).

## Data and code availability

All previously published scRNAseq data are downloaded and available on GEO per table 1. Upon publication, these data will be freely accessible. For the review process, the raw data can be accessed by GEO. DS18 - serie GSE280655, access token: **gzgrwmgmvfarvin**. DS19 - 22 GSE271016 access token: **ctwvmwsklretpwz**. DS23, 24 GSE271947, access token: **ylcpcmucdtslnch**. DS26 and striatal interneuron sequencing data could not be loaded onto GEO due to the US government shutdown, but will be loaded onto GEO at a later time. Meanwhile, they can be requested through the lead contact. Dev-SST-v2 can be accessed through a web interface https://shiny.gbiomed.kuleuven.be/AtlasLL/ or the final seurat object can also be requested through the lead contact. All customised analysis codes written in R, Python, Nextflow (Conda-environment) and Matlab used to generate Dev-SST-atlas and individual figure plots/ panels can be found on GitHub: https://github.com/Limlab-VIBCBD/Dev-SST-atlas.

## Extended data information

Extended data information includes 13 figures and 4 tables.

## Acknowledgement

We thank Boaz Levi and Jonathan Ting (Allen Institute) for providing the pAAV-eHGT-600m-sYFP2 plasmid, Zizhen Yao (Allen Institute) for early access to scRNAseq data from the developmental mouse visual cortex (Gao et al., 2024 preprint ^54^), and Marina Bershteyn (Neurona Therapeutics) for access to scRNAseq data from hPSC-derived MGE neurons xenotransplanted in mouse (Bershteyn et al., 2024 ^104^). We acknowledge the VIB-CBD expertise units - Single Cell Microfluidics, Genomics & Nucleomics, and Flow Cytometry - for their technical support. We are grateful to all members of the Lim lab, Marieke Verhagen, and Pierre Vanderhaeghen for discussions. This work was supported by the Research Foundation – Flanders (FWO) Junior Research Project (G057121N), FWO-Odysseus Award (G0E9121N), FWO/FNRS EOS (G0I7822N), and VIB Tech Watch grant (Techwatch - Vizgene) to L.L.; FWO Junior/Senior Research Projects (G074823N and G0A3Y24N) and International Foundation for Research in Paraplegia, Research Grant (P188) to AT.; National Institutes of Health grants R01NS121223 and RF1MH126719 to G.L.; F31NS118982 to J.X.D.; FWO PhD fellowship (1192822N) to E.M.

## Author Contributions

Conceptualization: M.L., F.F., G.B, & L.L.

Methodology: M.L., E.M. F.F., R.M., E.N., G.B., D.K., K.W., M.R., H.B., J.X.D. & L.L.

Investigation: M.L., E.M., F.F., R.M., G.B., H.B., K.W., M.R., J.X.D. & L.L. Analysis: M.L, E.M., F.F., D.K. & G.B.

Writing – Methods: F.F., M.L., G.B., R.M., E.M. & L.L.

Writing – Draft, Review, & Editing: M.L, F.F., G.B. & L.L Writing – Review & Editing: L.L.

Funding Acquisition: L.L., A.T. & G.L Resources: L.L., A.T., K.W. & G.L.

Supervision, L.L., A.T., K.W. & G.L.

