## Supplementary Figs for "Single-cell transcriptome of developing inhibitory neurons reveals expanding and contracting modes of diversification"

### EXTENDED DATA

#### Extended data Table 1, Extended data Figure 1-12

### Single-cell transcriptome of developing inhibitory neurons reveals expanding and contracting modes of diversification.

Minhui Liu,[1,2,6] Facundo Ferrero Restelli,[1,2,6] Elia Micoli,[1,2,6] Giulia Barbiera,[3] Rani Moors,[1,2] Evelien Nouboers, [1,2] Malou Reverendo,[1] Jessica Xinyun Du,[4] Hannah Bertels,[2] Dimitris Konstantopoulos,[3] Keimpe Wierd,[1] Aya Takeoka,[5] Giordano Lippi,[4] Lynette Lim[1,2,7]*

[1] VIB Center for Brain and Disease, 3000, Leuven, Belgium
[2] Department of Neurosciences, Katholieke Universiteit (KU) Leuven, 3000, Leuven, Belgium
[3] Genevia Technologies Oy, Tampere, Finland
[4] Department of Neuroscience, Scripps Research Institute, La Jolla, United States of America
[5] RIKEN Center for Brain Science, Saitama, 351-0198, Japan
[6] These authors contributed equally.

[7] Lead contact

| Extended Data Table 1: Mouse cortical developmental datasets | | | | | | | |
|---|---|---|---|---|---|---|---|
| **No.** | **Published Data** | **stages** | **Total cell** | **SST+ cell** | **Method** | **Location (if known)** | **Ref** |
| DS1 | GSE161605 | E12.5 | 4225 | 29 | Live cells dissociation | | 35 |
| DS2 | GSE211534 | E14.5 | 59537 | 586 | Nuclei dissociation | | 36 |
| DS3 | GSE161690 | E12.5, E14.5, E15.5, E16.5 & E18.5 | 10321 | 355 | Live cells dissociation | | 37 |
| DS4 | GSE184879 | E14.5 | 10931 | 148 | Live cells dissociation | | 38 |
| DS5 | GSE123335 | E14.5 and P0 | 18545 | 368 | Live cells dissociation | | 39 |
| DS6 | GSE153164 | E16.5, E18.5, P1, & P4 | 46282 | 3922 | Live cells dissociation | | 40 |
| DS7 | GSE104158 | E18.5 & P10 | 14971 | 5621 | Live cells dissociation + FACS enrichment | | 26 |
| DS8 | PRJNA637987 | E16.5 & E18.5 | 13760 | 824 | Live cells dissociation | | 41 |
| DS9 | GSE190940 | P8 | 1811 | 467 | Live cells dissociation | | 42 |
| DS10 | GSE181349 | E14.5 | 16892 | 0 | Live cells dissociation | | 43 |
| DS11 | GSM4195054 | P10 | 6313 | 346 | Live cells dissociation | | 44 |
| DS12 | GSM5014302, GSM5014304, GSM5014305, GSM5014306 | E13.5, P2 & P10 | 16948 | 4961 | For E13.5 & P2, Live cells dissociation For P10, Nuclei dissociation + FACS enrichment | | 45 |
| DS13 | GSE147247 | E18.5 | 12938 | 975 | Live cells dissociation | | 46 |
| DS14 | GSE188528 | E13.5 & E15.5 | 8740 | 232 | Live cells dissociation + FACS enrichment | | 47 |
| DS15 | GSE107122 | E13.5 & E15.5 | 10506 | 574 | Live cells dissociation | | 48 |
| DS16 | GSE93421 | E18.5 | 130000 0 | 7757 | Live cells dissociation | | 49 |
| DS17 | GSE217065, GSE235619 | E16.5, P1, & P5 | 23801 | 19874 | For E16.5, P1, P5-1, Live cells dissociation + FACS enrichment For P5-2, Fixed cells + FACS enrichment | Live cells - KCL UK; Fixed cell - VIB-CBD, Leuven, BE | 23 |
| DS25 | GSM8409508 | P2 & P7 | 5919 | 2469 | Nuclei dissociation + FACS enrichment | Harvard University, Boston USA | 68 |
| | **new datasets** | **stages** | **Total cell** | **SST+ cell** | **Method** | **Location (if known)** | **Ref** |
| DS18 | GSE280655 | E18.5 | 8015 | 2431 | Live cells dissociation + FACS enrichment | Scripps, San Diego, USA | this study |
| DS19 | GSM8367247 | P5-1 | 8722 | 5316 | Live cells dissociation + FACS enrichment | VIB-CBD, Leuven, BE | this study |
| DS20 | GSM8367248 | P5-2 | 10431 | 6546 | Live cells dissociation + FACS enrichment | VIB-CBD, Leuven, BE | this study |
| DS21 | GSM8367249 | E16.5-1 | 7882 | 5301 | Live cells dissociation + FACS enrichment | VIB-CBD, Leuven, BE | this study |
| DS22 | GSM8367250 | P1-1 | 5820 | 4200 | Live cells dissociation + FACS enrichment | VIB-CBD, Leuven, BE | this study |
| DS23 | GSM8389469 | P1-2 | 10474 | 1763 | Fixed cells + FACS enrichment | VIB-CBD, Leuven, BE | this study |
| DS24 | GSM8389467 | P1-3 | 18417 | 9758 | Live cells dissociation + FACS enrichment | VIB-CBD, Leuven, BE | this study |
| DS26 | To be uploaded | E16.5-2 | 7552 | 2563 | Live cells dissociation + FACS enrichment | VIB-CBD, Leuven, BE | this study |

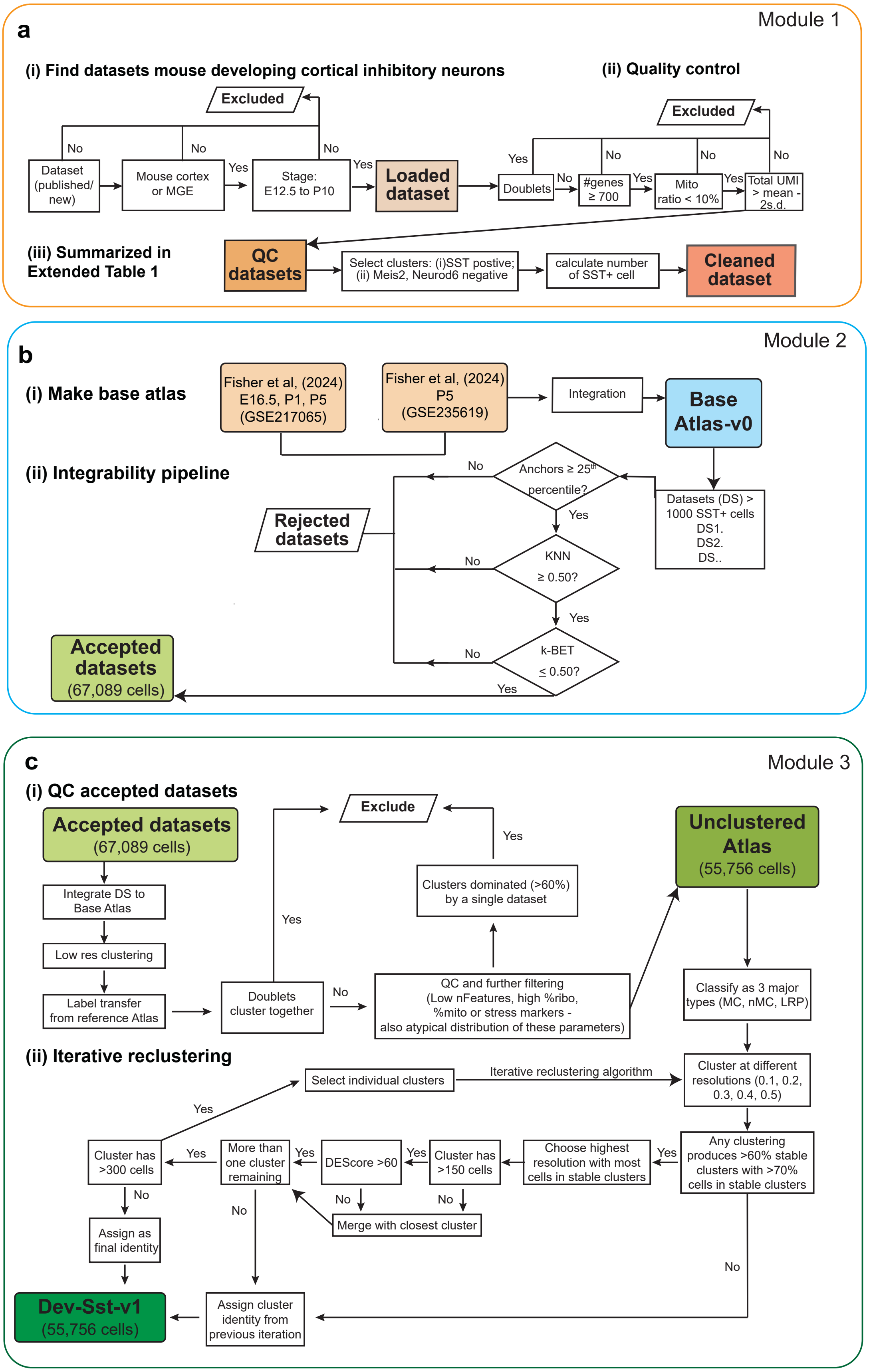


**Extended data Fig.1. Computation pipeline for the generation of Dev-Sst-v1.**
(**a**) Module 1 flowchart outlining the filtering criteria, QC modules and steps for isolating high-quality cortical Sst+ cells.

(**b**) Module 2 flowchart illustrating the integrable testing using Atlas-v0 as the reference atlas.
(**c**) Module 3 flowchart detailing the additional quality control steps applied to Atlas-v1, following the re-clustering algorithm used to generate Dev-Sst-v1 from Atlas-v1.

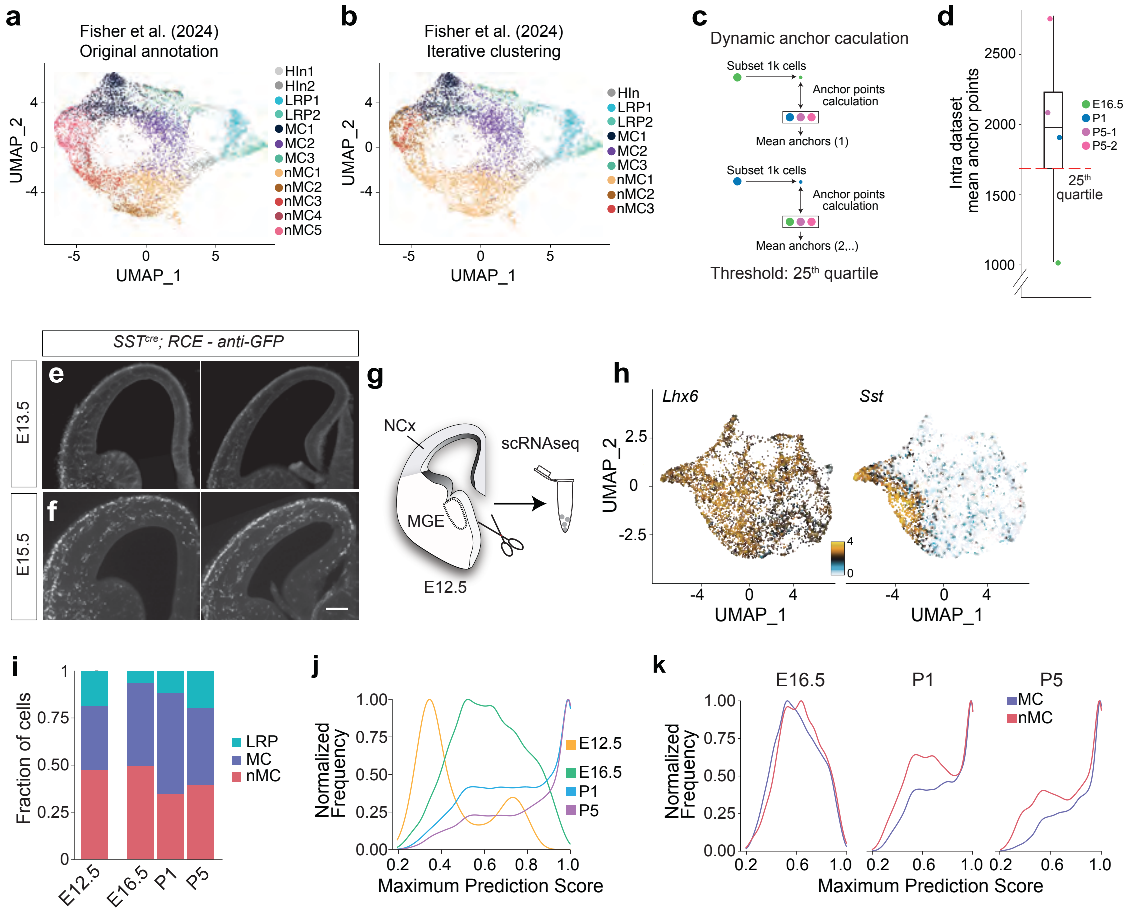


**Extended Data Figure 2. Validation of clustering parameters and developmental constraints on cross-dataset mapping.**

(**a, b**) Application of the clustering parameters used in this study (see Methods) to the original datasets from Fisher et al. (2024) reproduced a comparable number of clusters relative to the published annotation.

(**c, d**) For each of the four reference datasets, 1,000 cells were randomly subsampled and used to compute mean anchor points against the remaining datasets. Anchor distributions ranged from 1,021 to 2,775 anchors across samples.

(**e, f**) Coronal sections of *SST$^{cre}$;RCE* mice at (e) E13.5 and (f) E15.5 showing only a small number of SST$^{+}$ interneurons had reached the cortex at E13.5.

**(g)** Schema on solation of MGE tissue at E12.5 captured newly generated post-mitotic SST$^{+}$ interneurons.

**(h)** UMAP of E12.5 scRNAseq with expression of Sst and Lhx6.

**(i)** Fraction of cells could be mapped broadly to MC, nMC, and LRP groups at various timepoints.

**(j)** Mapping probabilities for E12.5 cells were markedly reduced relative to samples from E16.5 and older.

**(k)** After E16.5, MC and nMC clusters showed no detectable differences in mapping probability.

a

| Dataset | Stages | Spinal cord neuronal nuclei | Mean anchors | k-BET test | Fraction within k-NN |
|---|---|---|---|---|---|
| I2 | P90 | 12813 | 3574.15 | 0.34 | 0.86 |
| I3 | P90 | 13064 | 4261.14 | 0.08 | 0.99 |
| I4 | P87 | 19600 | 3859.25 | 0.47 | 0.82 |
| N1 | P66 | 16855 | 4013.18 | 0.59 | 0.71 |
| N2 | P66 | 9908 | 3984.10 | 0.27 | 0.86 |
| N3 | P66 | 16757 | 3541.65 | 0.66 | 0.70 |
| N4 | P66 | 9752 | 3835.30 | 0.36 | 0.94 |
| A1 | P93 | 14727 | 4347.60 | 0.23 | 0.94 |
| A2 | P89 | 10864 | 3897.73 | 0.44 | 0.81 |
| A3 | P87 | 10948 | 4151.09 | 0.11 | 0.98 |
| A4 | P89 | 5599 | 3799.00 | 0.23 | 0.90 |
| Af1 | P93 | 13394 | 4291.21 | 0.19 | 0.94 |
| Af2 | P110 | 16273 | 3285.35 | 0.69 | 0.60 |

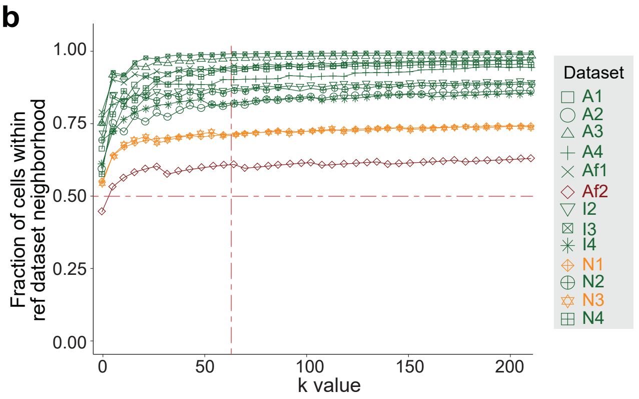


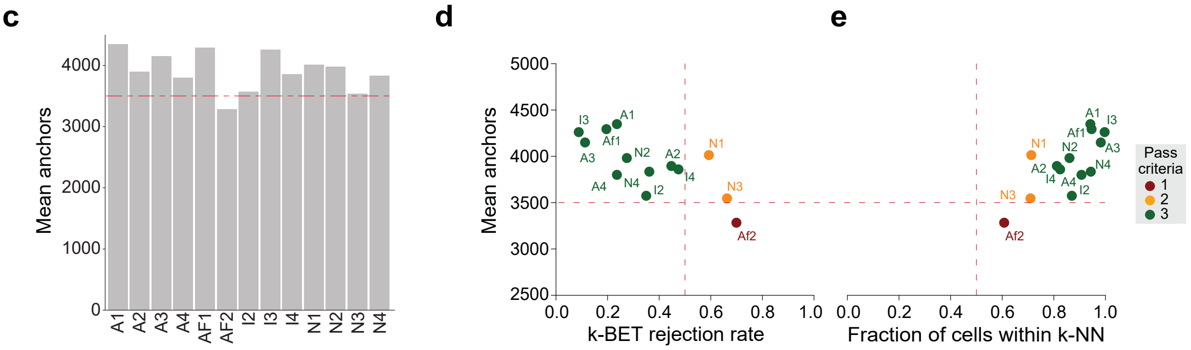


**Extended Data Fig. 3. Appling Module 2 to assess adult spinal cord single-nuclei RNAseq datasets.**

(**a**) Table summarizing the developmental stages, number of nuclei, anchor points, k-BET rejection rates, and k-NN values for 13 different spinal cord snRNAseq datasets.

(**b**) Fraction of cells in each dataset within reference neighborhoods with increasing k value. The dashed line (k = 60) indicates the acceptance threshold of >50%.

(**c**) Bar plot of anchor points calculated using Seurat's Canonical Correlation Analysis (CCA) integration function between each dataset and reference.

(**d**) Dot plot illustrating the relationship between k-BET rejection rates and

(**e**) fraction of cells included in k-NN (k=60) versus the number of anchor points. Datasets passing the three integrability criteria (green): for these series of experiments, the mean anchor points >15000, fraction within k-NN >60%, and k-BET rejection rate <0.6.

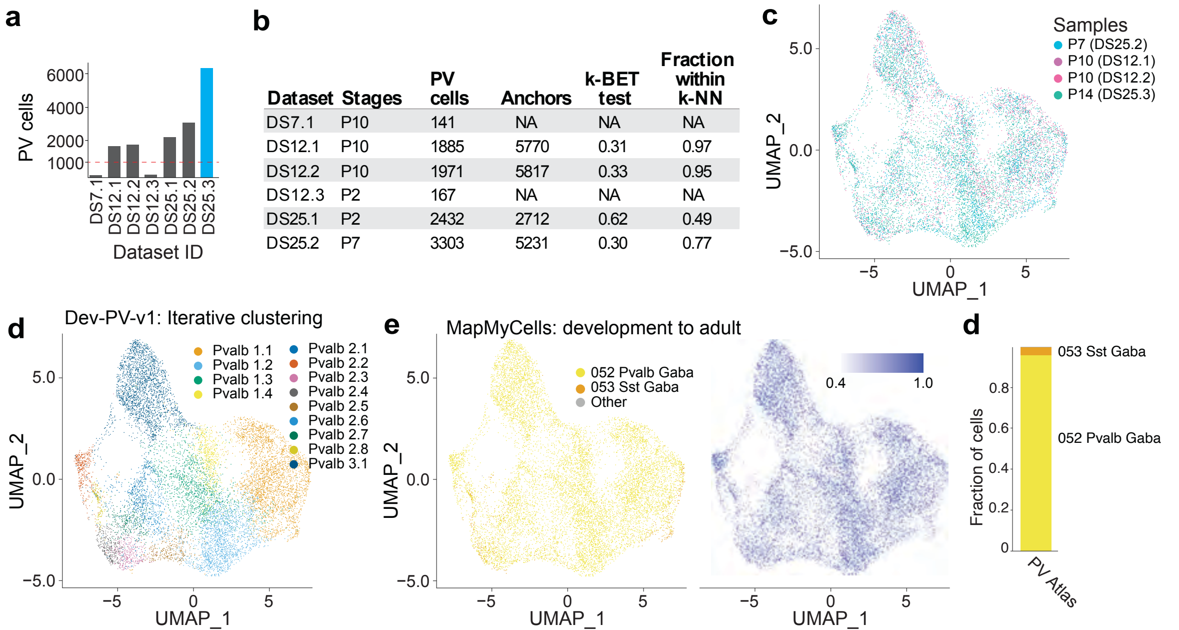


| Dataset | Stages | PV cells | Anchors | k-BET test | Fraction within k-NN |
|---|---|---|---|---|---|
| DS7.1 | P10 | 141 | NA | NA | NA |
| DS12.1 | P10 | 1885 | 5770 | 0.31 | 0.97 |
| DS12.2 | P10 | 1971 | 5817 | 0.33 | 0.95 |
| DS12.3 | P2 | 167 | NA | NA | NA |
| DS25.1 | P2 | 2432 | 2712 | 0.62 | 0.49 |
| DS25.2 | P7 | 3303 | 5231 | 0.30 | 0.77 |

**Extended Data Figure 4. Construction of a developmental atlas of cortical PV⁺ interneurons using cross-dataset integration criteria.**

**(a)** Number of PV⁺ interneurons identified across six publicly available datasets (DS7.1, DS12.1–12.3, DS25.1–25.2). The dataset with the largest number of PV⁺ cells was selected as the base reference for atlas construction.

**(b)** Evaluation of datasets using three integration metrics—anchor point number, fraction of cells within k-nearest neighbors (k = 60), and the k-BET batch-mixing test.

**(c)** Four datasets that passed all inclusion criteria were integrated to construct the developmental PV atlas.

**(d)** UMAP labeling iterative clustering of the integrated dataset produced the Dev-PV-v1 atlas, consisting of multiple transcriptionally defined Pvalb-lineage clusters.

**(e, f)** Using Mapmycells, >85% of cells within Dev-PV-v1 mapped to the adult 052 Pvalb Gaba reference cluster with prediction probability >0.9, demonstrating high cross-stage concordance.

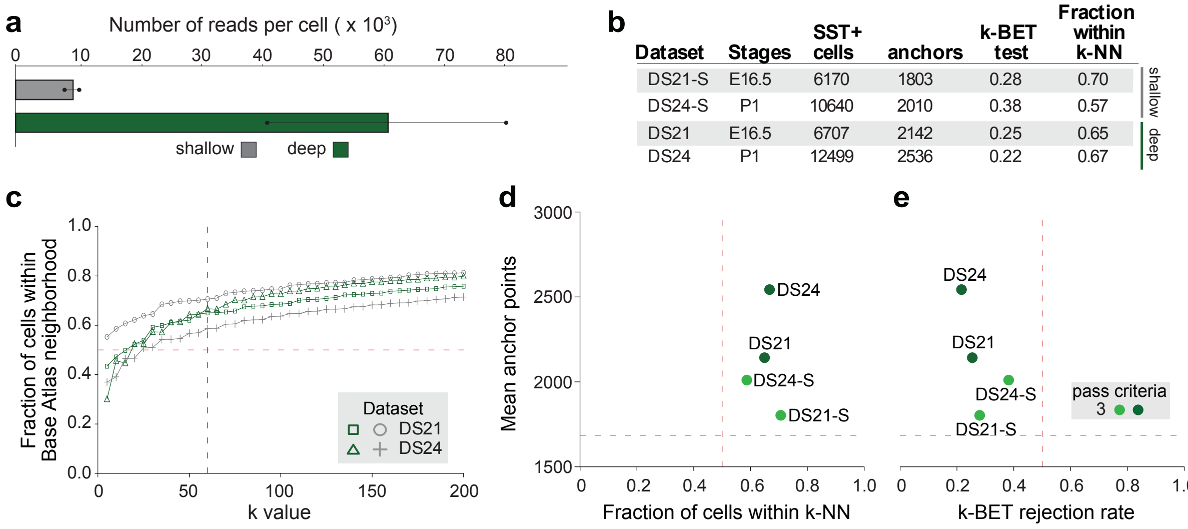


**Extended Data Figure 5. Integrability of the same dataset with variations in sequencing depth.**
(**a**) Comparison of sequencing depth between shallow and deep sequencing datasets DS21 and DS24. Shallow sequencing targeted 5,000 reads per cell (mean achieved: 8,227 ± 794 reads per cell), while deep sequencing targeted 50,000 reads per cell (mean achieved: 61,384 ± 20,484 reads per cell).
(**b**) Normalized mean anchor points for DS21 and DS24 under shallow (S) and deep sequencing protocols. While DS21-S showed a slight reduction in anchor points below the acceptance threshold of 1686 (horizontal dashed line), DS24-S maintained integrability.
(**c**) Fraction of cells in shallow- and deep-sequenced datasets within Dev-SST-v0 neighborhoods with increasing k value. Both shallow and deep sequencing protocols met the acceptance threshold of >50% (k=60) (dashed line).
(**d**, **e**) Dot plot of (d) fraction cells within k-NN (k=60) and (**e**) k-BET rejection rates plotted against anchor points for shallow and deep sequencing. Datasets of either deep and shallow sequencing depth that passed most integrability criteria (in green) have mean anchor points >2000, fraction within k-NN >50%, and k-BET rejection rate <0.5.

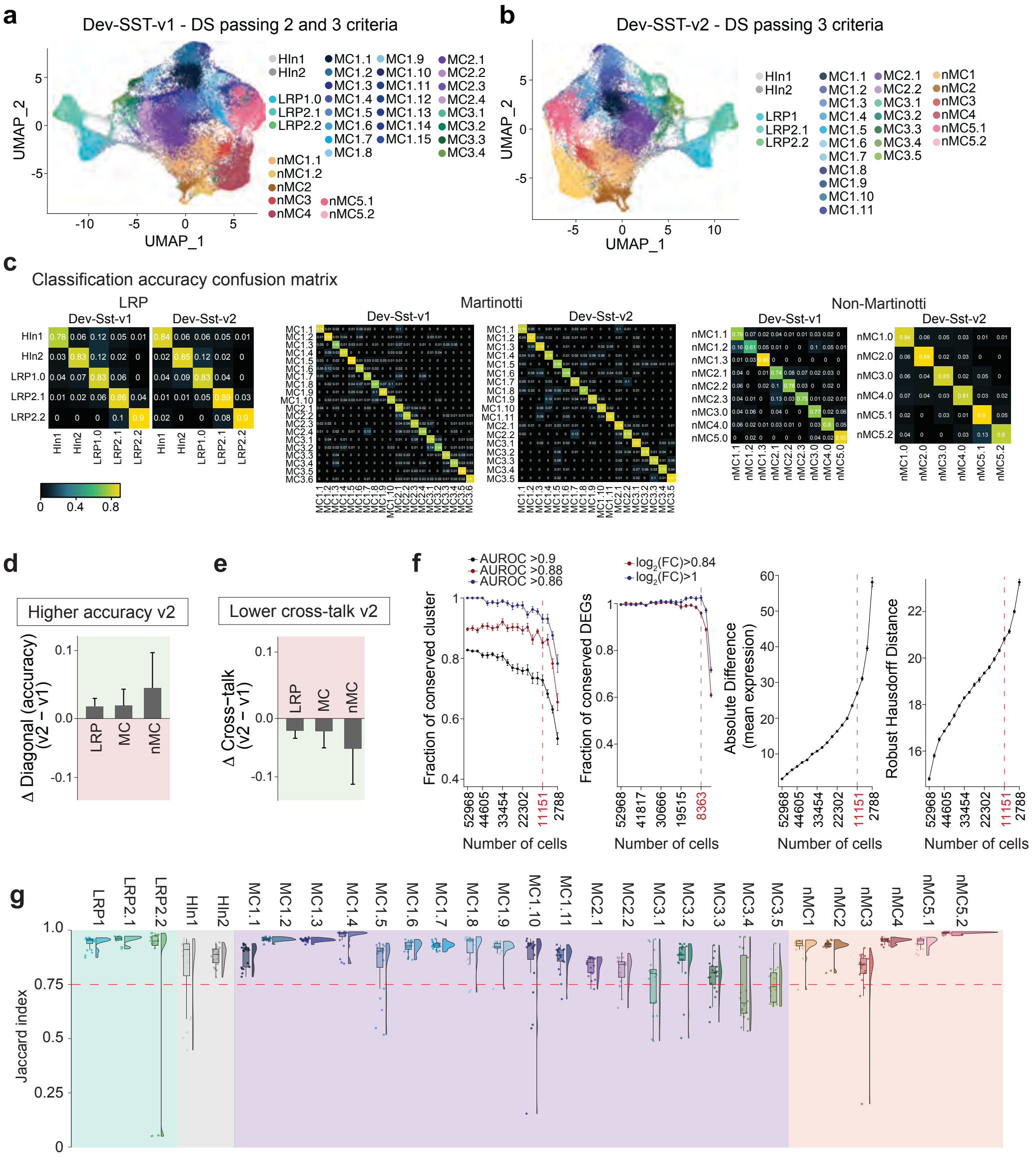

**Extended Data Figure 6. Construction, comparison, and validation of two developmental SST⁺ interneuron atlases (Dev-SST-v1 and Dev-SST-v2).**

**(a, b)** UMAP of SST⁺ neurons integrated datasets that passed either ≥2 criteria (**a,** Dev-SST-v1) or all 3 criteria (**b,** Dev-SST-v2) in Module 2 using Seurat CCA. The final Dev-SST-v1 and Dev-SST-v2 atlases contained 63,572 cells (stages E16.5, E18.5, P1, P2, P5, P7, P10) and 59,658 cells (stages E16.5, E18.5, P1, P2, P5), respectively.

**(c)** Confusion matrices derived from random forest classification summarizing how true cell identities in each atlas version were assigned to predicted types. Diagonal entries represent correct classifications (accuracy), while off-diagonal entries capture misclassifications (cross-talk).

**(d, e)** Per-class comparisons between versions (Δ = v2 − v1) for diagonal accuracy **(d)** and cross-talk **(e)** showed that Dev-SST-v2 consistently improved or maintained accuracy and reduced cross-talk across Martinotti (MC), non-Martinotti (nMC), and LRP populations.

**(f)** Saturation analysis assessing sampling sufficiency across 2,560–48,656 subsampled cells. For each subsample, four metrics were computed: (i) fraction of conserved clusters, (ii) fraction of conserved differentially expressed genes, (iii) absolute difference in mean expression, and (iv) the

robust Hausdorff distance. Inflection points from second-derivative analyses identified saturation at 8,639–11,151 cells. All major developmental stages (E16.5, E18.5/P1, P5) exceed this threshold.
**(g)** Violin plot of Jaccard index of all clusters in Dev-SST-v2. The cutoff of 0.75, red dashed line, was used for cluster retention.

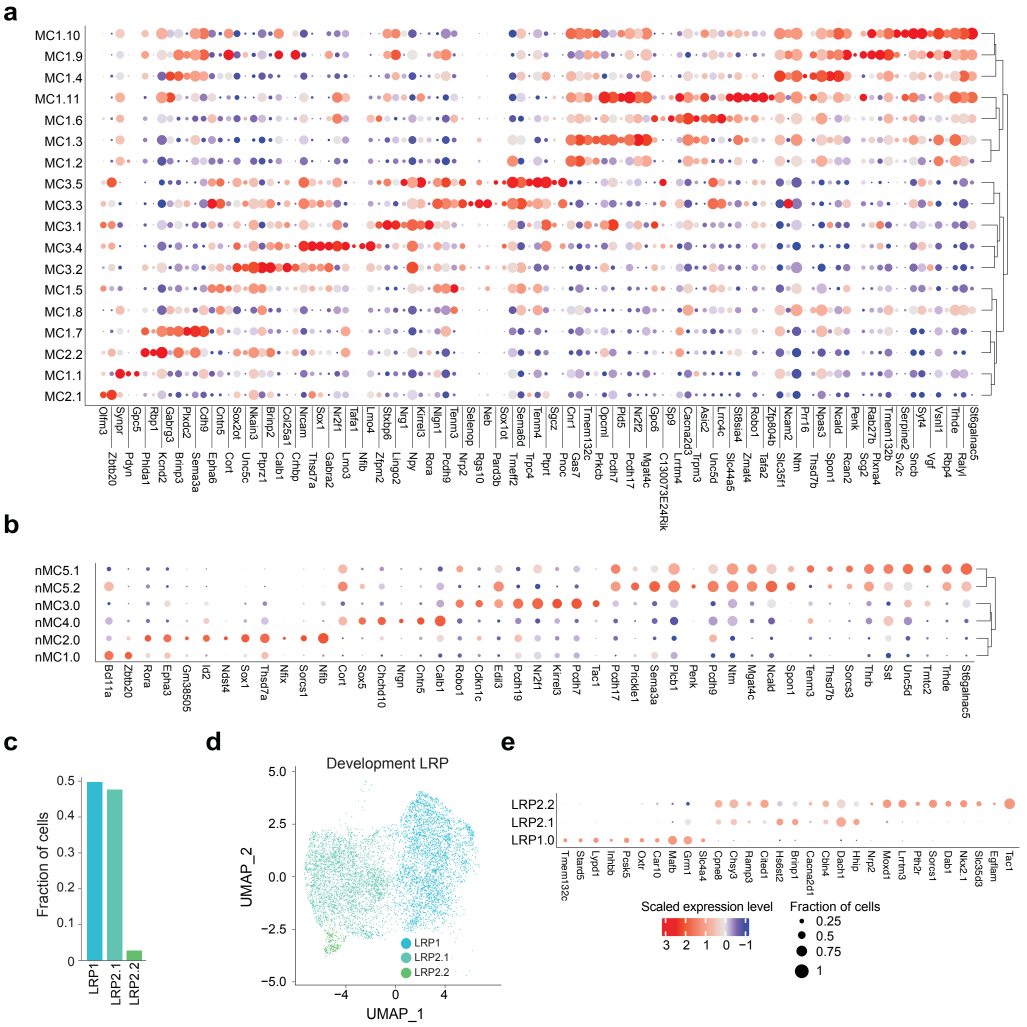


**Extended Data Fig. 7. Gene expression differences among distinct clusters within Dev-Sst-v1.**
(**a**, **b**) Bubble plots of marker genes in clusters of (**d**) MCs (**e**) and nMCs.
(**c**) Barplot showing the relative fraction of LRP types within the LRP population. LRP2.2 < 2%.
(**d**) UMAP projections of LRP types from Dev-Sst-v1.
(**e**) Bubble plots showing marker gene expression of each LRP. Note LRP2.2 exhibits high expression of Nkx2.1, a marker for striatal SST+ interneuron.

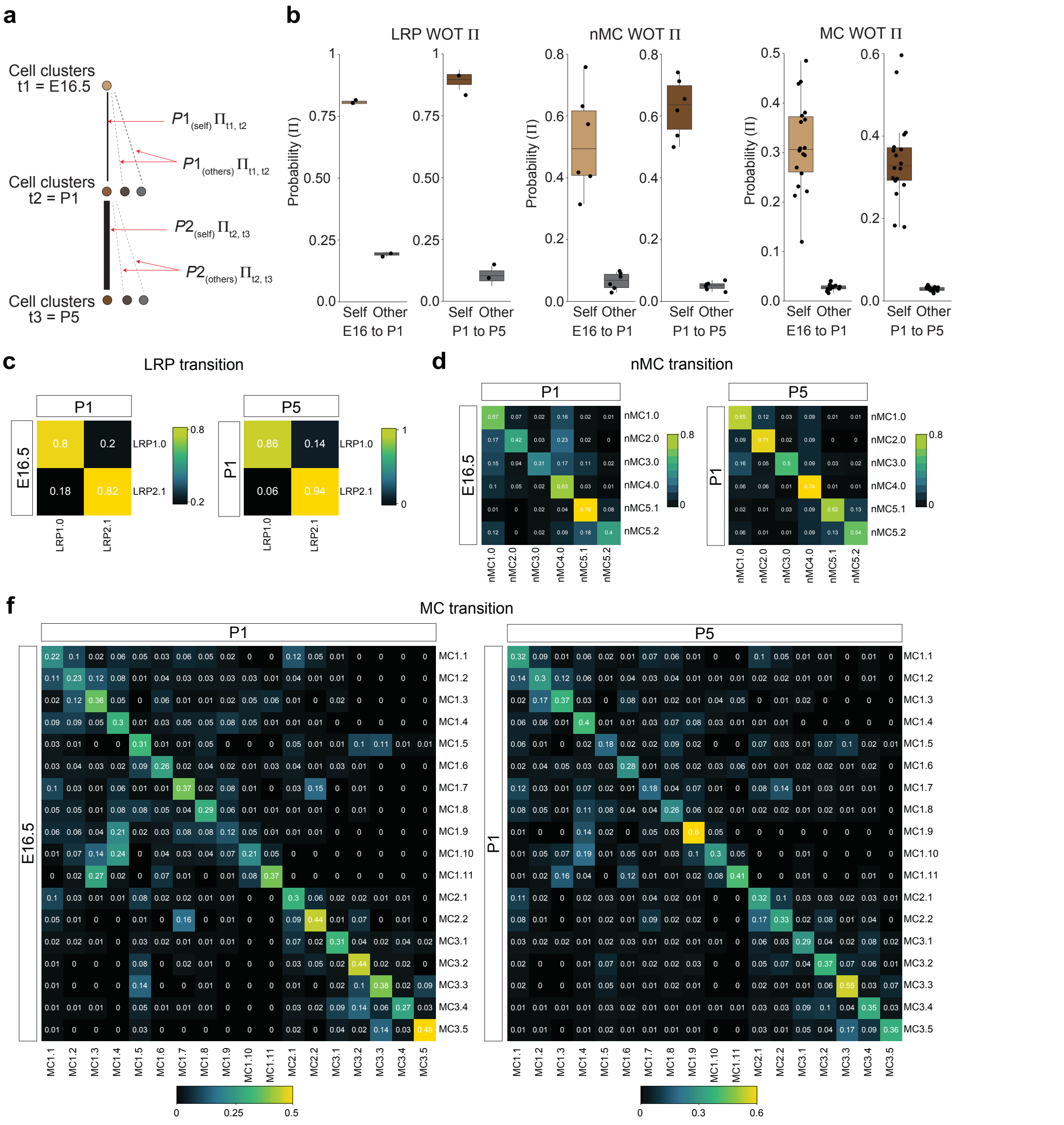


**Extended Data Figure 8. Waddington Optimal Transport analysis validates developmental transitions inferred from CCA-based integration.**

**(a)** Schema of Waddington Optimal Transport (WOT) probabilistic transitions between successive developmental stages (E16.5 → P1 → P5).

(**b)** Strong enrichment of diagonal couplings relative to off-diagonal transitions was observed for LRP, Martinotti (MC), and non-Martinotti (nMC) cells, indicating that cells predominantly transition into the expected lineage-matched clusters across time.

(**c – f**) Confusion matrix of WOT probability for E16 to P1 and P1 to P5 transition for LPR **(c)**, nMC **(d)**, and MC **(f)**.

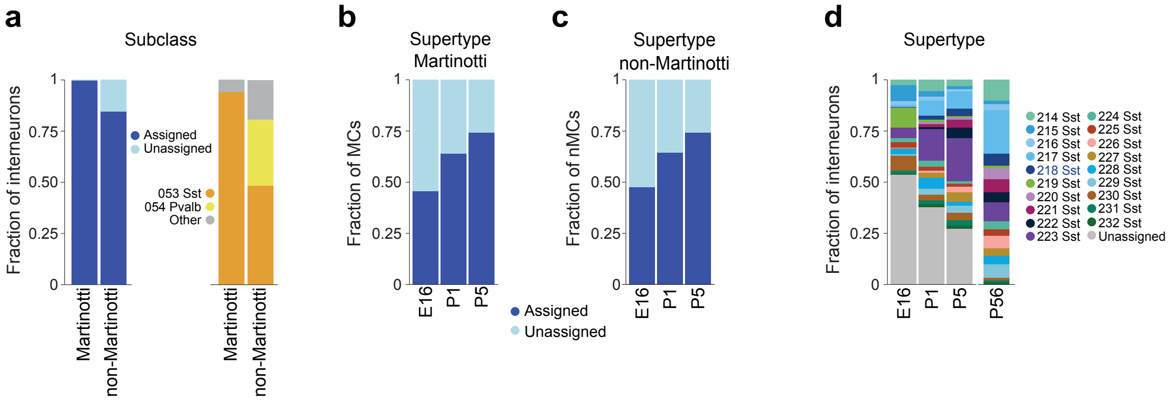


**Extended Data Fig. 9. Assignment of developmental MCs and nMCs to adult subclasses and supertype using MapMyCells.**

(**a**) Left: stacked bar plot representing fraction of MCs and nMCs that are assigned to adult subclasses. Assignments with >0.9 or <0.9 probability are annotated as “asssigned” or “unassigned” respectively. Right: assignment of different subclasses with probability >0.9 .

(**b**, **c**) Stacked bar plot representing fraction of (**b**) MCs and (**c**) nMCs at different stages that are assigned with a high probability (>0.9) to adult supertypes.

(**d**) Stacked bar plot representing developmental SST interneuron assigned to each supertype with a high probability (>0.9).

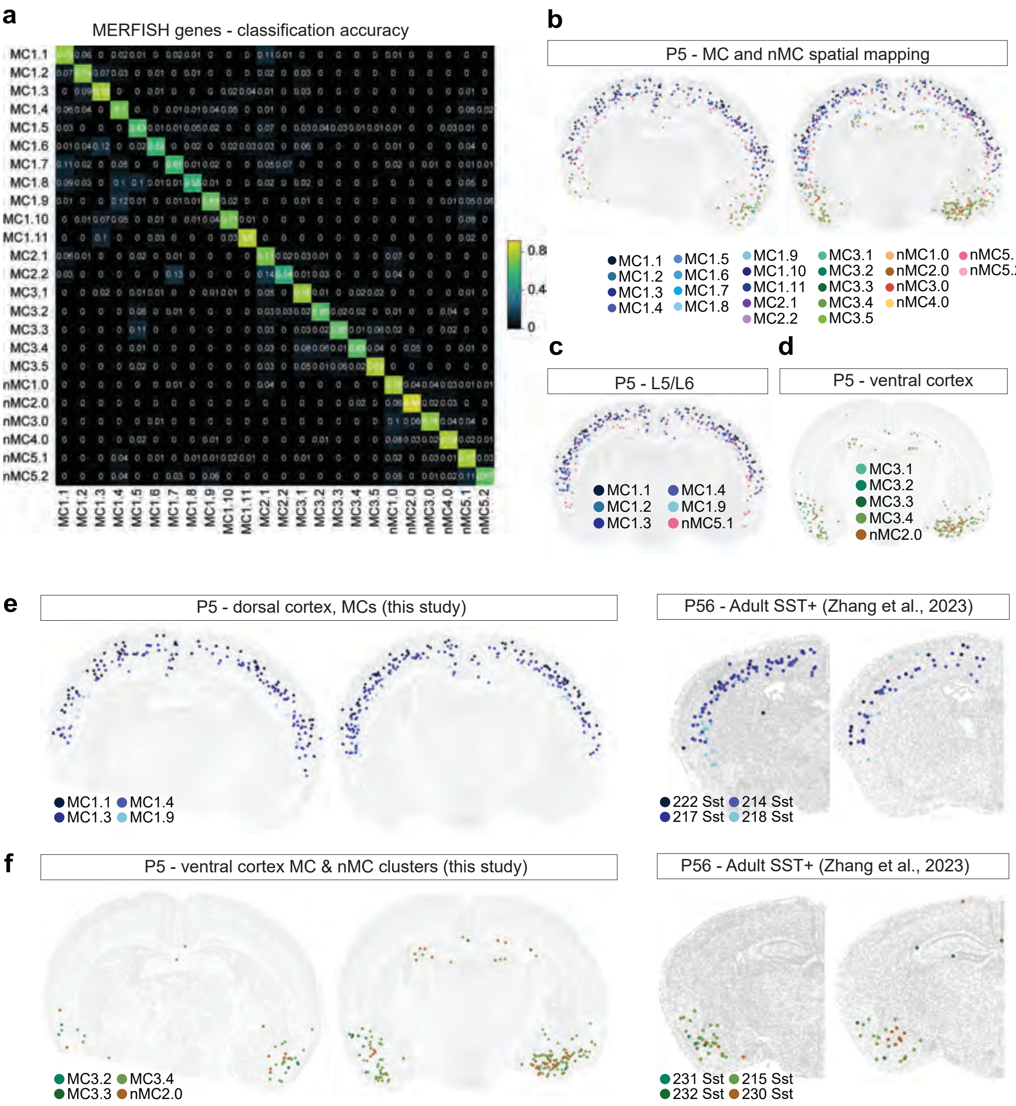


**Extended Data Figure 10. P5 MERFISH spatial validation of developmental SST⁺ interneuron clusters.**

**(a)** Confusion matrix on classifying accuracy of customized probe set of ~500 genes (Extended Data Table 2) used in MERFISH. This gene set is optimized to classify Dev-SST-v2 clusters with high accuracy.

**(b)** Spatial cell type assignment using MERFISH. Most developmental clusters exhibited distinct and regionally enriched spatial distributions across the P5 cortex.

**(c)** Six clusters were confined to infragranular layers (L5/L6).

**(d)** Five clusters localized predominantly to ventral cortical regions, including insular and piriform cortices.

**(e, f)** To assess whether the spatial organization of developmental clusters resembles that of their adult counterparts, we compared P5 MERFISH data with adult SST⁺ MERFISH from Zhang *et al.* (2023). Eight developmental clusters—four infragranular and four ventral—were examined alongside their one-to-one adult Sst supertypes. All pairs displayed highly similar regional localization patterns in both developmental and adult tissue.

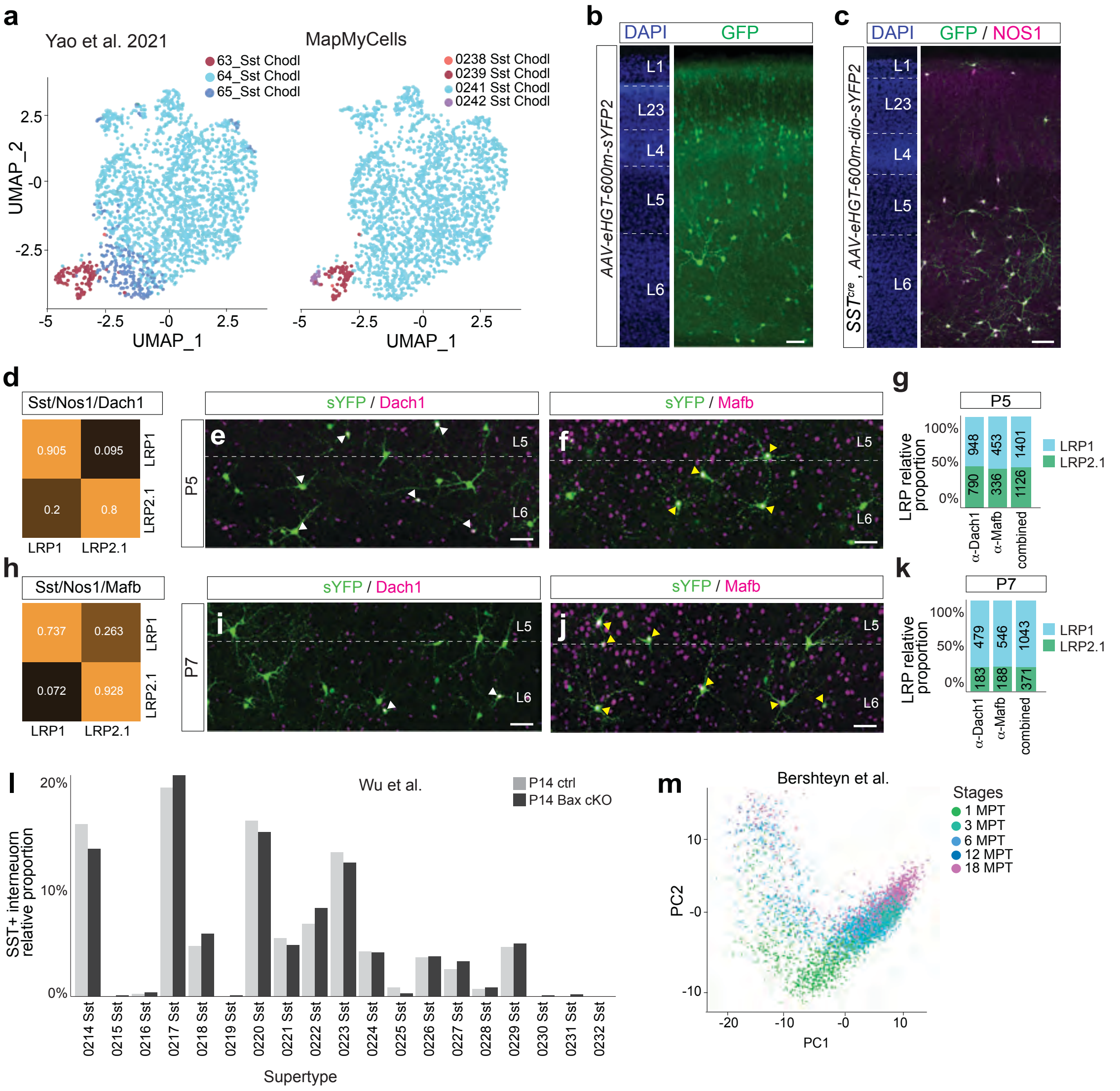


**Extended Data Fig. 6. Characterisation of LRP1 and LRP2.1**

(**a**) UMAP plot of Sst-Chodl clusters (LRP) found in Yao et al., 2021 using annotation in taht paper (left) and annotation assigned by MapMyCells (right). 63_Sst_Chodl cells are predominately assigned as supertype 0239 Sst Chodl.

(**b**) P6 cortical tissues injected with AAV-eGHT-600m-sYFP2 at P0 and staining with GFP

(**c**) P6 Sstcre cortical tissues injected with AAV-eGHT-dio-h600-sYFP2 at P0 and stained with GFP and NOS1. Scale bar = 100µm

(**d**, **h**) confusion matrix calculating accuracy of distinguishing LRP1 and LRP2.1 in Dev-Sst-v1 using 3 genes (d) *Sst/Nos1/Dach1* or (h) *Sst/Nos1/MafB*

(**e, f, i, j**) co-immunostaining of (**e, i**) GFP/Dach1 or (**f, j**) GFP/MafB with quantifying relative abundance of LRP1 and LRP2.1 in Sstcre using mice injected AAV2,9-eGHT-dio-600m-sYFP2 at P0 and immunostained at (**e, f**) P5 or (**i, j**) at P7. Yellow and white arrow heads denote LRP1 and LRP2.1 respectively.

(**g, k**) Barplots represent quantification of LRP1 and LRP2.1 assessed by staining with Dach1, Mafb, or combined - mean result of both methods at (**g**) P5 and (**k**) P7. The numbers of the bars

represent the cell number quantified from each condition, n = 3 mice per time point. Both methods give similar results.
(**l**) Relative abundance of 19 SST+ interneuron supertypes at P14 in control and Bax cKO mice, datasets reanalyzed from Wu et al, (2024 preprint [68]). No significant differences in the relative abundance of these supertypes were observed.
(**m**) PCA plots of hLRPs from Bershteyn et al., (2024 preprint [104]) labeled by stages post transplantation.

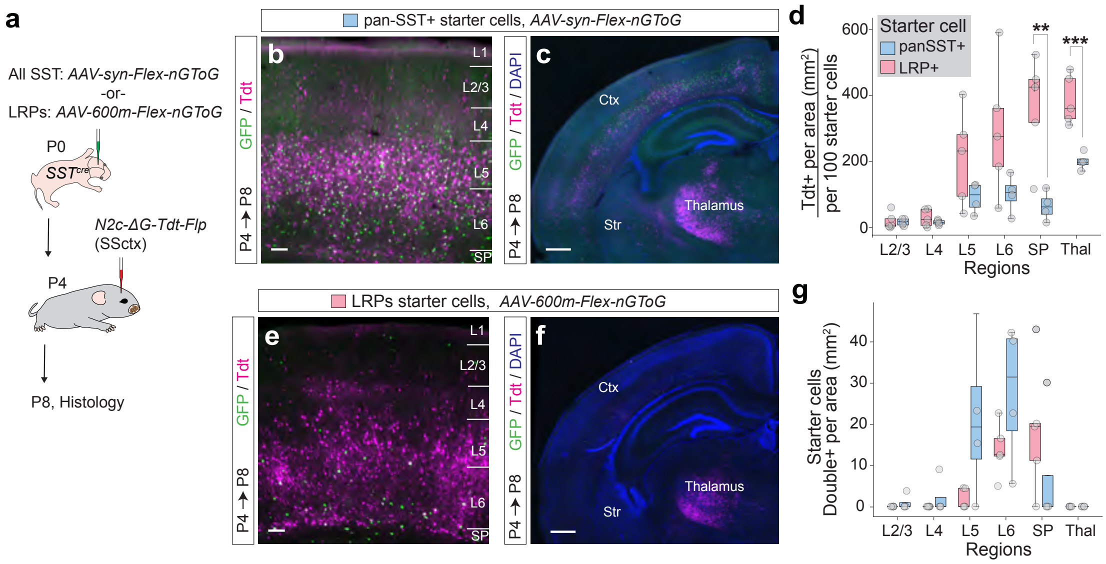


**Extended Data Figure 12. Rabies-based monosynaptic tracing reveals that LRPs receive strong thalamic and subplate inputs during early postnatal development.**

**(a)** Experimental strategy. *SST$^{Cre}$* mice were injected at P0 with Cre-dependent AAV helper viruses encoding GFP, TVA, and optimized rabies G protein. A synapsin-promoter helper virus (AAV2/9-syn-Flex-nGToG) labeled all SST$^+$ cells (pan-SST$^+$), whereas an LPR-specific enhancer helper virus (AAV2/9-600m-Flex-nGToG) selectively labeled LPRs. At P4, N2c-ΔG-Tdt-Flp rabies virus was injected into somatosensory cortex, and histology was performed at P8. Starter cells were defined as TdTomato$^+$/GFP$^+$ double-positive neurons.

**(b, c, e, f )** In mice labeled with the synapsin-promoter helper virus (b, c) or LPR-specific helper virus (e, f). Both conditions extensive TdTomato$^+$ presynaptic neurons were detected in thalamus and deep cortical layers

**(d)** Quantification of presynaptic inputs normalized to 100 starter cells showed robust labeling across cortical and thalamic regions for pan-SST$^+$ & LPR starter populations (**$p < 0.01$) and thalamic (***$p < 0.001$).

**(g)** Mean starter cell density of SST$^+$ cells from the synapsin-promoter helper virus or LPR-specific helper.
